# Systematic discovery of a topical bacterial consortium that targets *Staphylococcus aureus* to treat atopic dermatitis

**DOI:** 10.64898/2026.02.13.705787

**Authors:** Emily L. Bean, Bernardo Cervantes, Keaton Armentrout, A.H. Hassaballah, Omobukola Solebo, Heidi A. Arjes, Eliza A. Zalis, Alexander Lyons, Denny Chin, Peter Lio, Kevin D. Litcofsky, Cheri M. Ackerman Araromi, Jared Kehe

## Abstract

Atopic dermatitis (AD) flares are frequently accompanied by *Staphylococcus aureus* overgrowth and activation of quorum sensing-regulated virulence pathways that amplify inflammation and barrier dysfunction. Because commensal members of the skin microbiome can inhibit *S. aureus* colonization and virulence, we hypothesized that a consortium sourced from healthy human skin could therapeutically target *S. aureus* and ameliorate AD. We curated 180 skin-derived bacterial strains and used kChip, an ultrahigh-throughput coculture platform, to profile *S. aureus*’ physiological response to over four million combinations consisting of two, three, or seven cocultured strains. This screening identified Ensemble No.2 (ENS-002), a three-strain consortium that strongly suppressed *S. aureus* growth and virulence in follow-up microtiter assays and a Reconstructed Human Epidermis *S. aureus* Activity (RHESA) model. ENS-002 is now undergoing development as a topical live biotherapeutic product (LBP) treatment for AD.

## Introduction

Atopic dermatitis (AD) is a chronic inflammatory skin condition characterized by intense pruritus, erythematous lesions, and skin barrier dysfunction. Although AD etiology is complex and multifactorial, extensive evidence links *S. aureus* colonization to AD pathogenesis, with *S. aureus* blooms and a concurrent decrease in microbial diversity frequently preceding disease flares^1,2^. Between 70 and 93% of patients with AD have skin lesions colonized by *S. aureus*^3–5^, with the prevalence of *S. aureus* carriage on lesional skin increasing with AD severity^5,6^.

AD is sustained by a reinforcing cycle between epidermal inflammation and *S. aureus* colonization, in which each process potentiates the other. *S. aureus* activates quorum sensing-regulated virulence programs that damage the epidermal barrier, provoke itch, and amplify the type 2 inflammatory response (for reviews, see ^7–13^). In turn, type 2 inflammation increases extracellular matrix protein production and decreases antimicrobial peptide (AMP) expression^14–19^, creating favorable conditions for *S. aureus* adhesion and expansion.

Successful AD treatments can interrupt the mutual reinforcement of inflammation and *S. aureus* overgrowth but have limitations for long-term management of mild-to-moderate disease. Biologic agents that target type 2 cytokine signalling, such as dupilumab, can reduce *S. aureus* colonization indirectly by dampening the inflammatory environment that favors its growth^20^. Although systemic immunomodulation may be suitable for long-term use, patients with a body surface area (BSA) involvement <10% and low severity scores may not qualify for treatment. Antibiotics that directly suppress *S. aureus* can improve AD^21–26^ but have a short-term impact, as they disrupt the skin microbiome, promote antibiotic resistance, and in some cases allow *S. aureus* to re-colonize the vacant skin niche after treatment cessation^22,27^. Accordingly, there remains a need for therapies suitable for safe, sustained use that durably suppress *S. aureus* overgrowth and virulence in mild-to-moderate AD^28–30^.

Protective *S. aureus*-suppressing mechanisms innate to the healthy skin microbiome present a promising therapeutic avenue in AD management. Commensal microbes can work cooperatively to inhibit *S. aureus* colonization and overgrowth^31–33^. Several live biotherapeutic products (LBPs) have been investigated to reestablish microbial control over *S. aureus*^1,14,31–39^, including a strain of *Staphylococcus hominis* that showed clinical benefit among patients for whom *S. aureus* burden decreased during treatment^37^.

We set out to systematically identify an LBP containing multiple therapeutic species that could robustly pacify *S. aureus* and therefore treat AD. These strains could have complementary or redundant mechanisms that impact *S. aureus* metabolic activity, viability, and virulence, and including more strains would provide more opportunities for successful revival and activity across diverse skin environments.

Here, we present the discovery of “Ensemble No.2” (ENS-002), a topical three-strain LBP designed to treat AD by suppressing *S. aureus* growth and virulence. To identify these strains, we screened tens of thousands of pairwise and higher-order combinations of skin commensals for their ability to suppress *S. aureus* metabolic activity and virulence factor expression. As this screen would not be feasible with current standard laboratory techniques, we employed kChip, an ultrahigh-throughput combinatorial screening platform^40,41^. With kChip, we screened over 4 million defined microbial communities for their ability to influence expression of multiple *S. aureus* genes implicated in AD. The identified members of ENS-002—strains ii283-N (*Bacillus* “Strain N”), ji77-T (*Lysinibacillus* “Strain T”), and ii332-X (*Bacillus* “Strain X”)—together robustly suppressed *S. aureus* across kChip communities, standard broth coculture assays, and reconstructed human epidermis (RHE) models.

## Results

### A diverse and manufacturable set of skin strains was curated for screening

To develop an LBP suitable for treating AD, we aimed to identify a combination of bacterial strains that was safe and tolerable on human skin, culturable in a manner amenable to manufacturing, and able to robustly suppress *S. aureus* virulence. To this end, we assembled a skin-derived bacterial biobank and applied an iterative downselection screening process to identify candidate strains for therapeutic development (Fig. 1a).

**Fig. 1.**
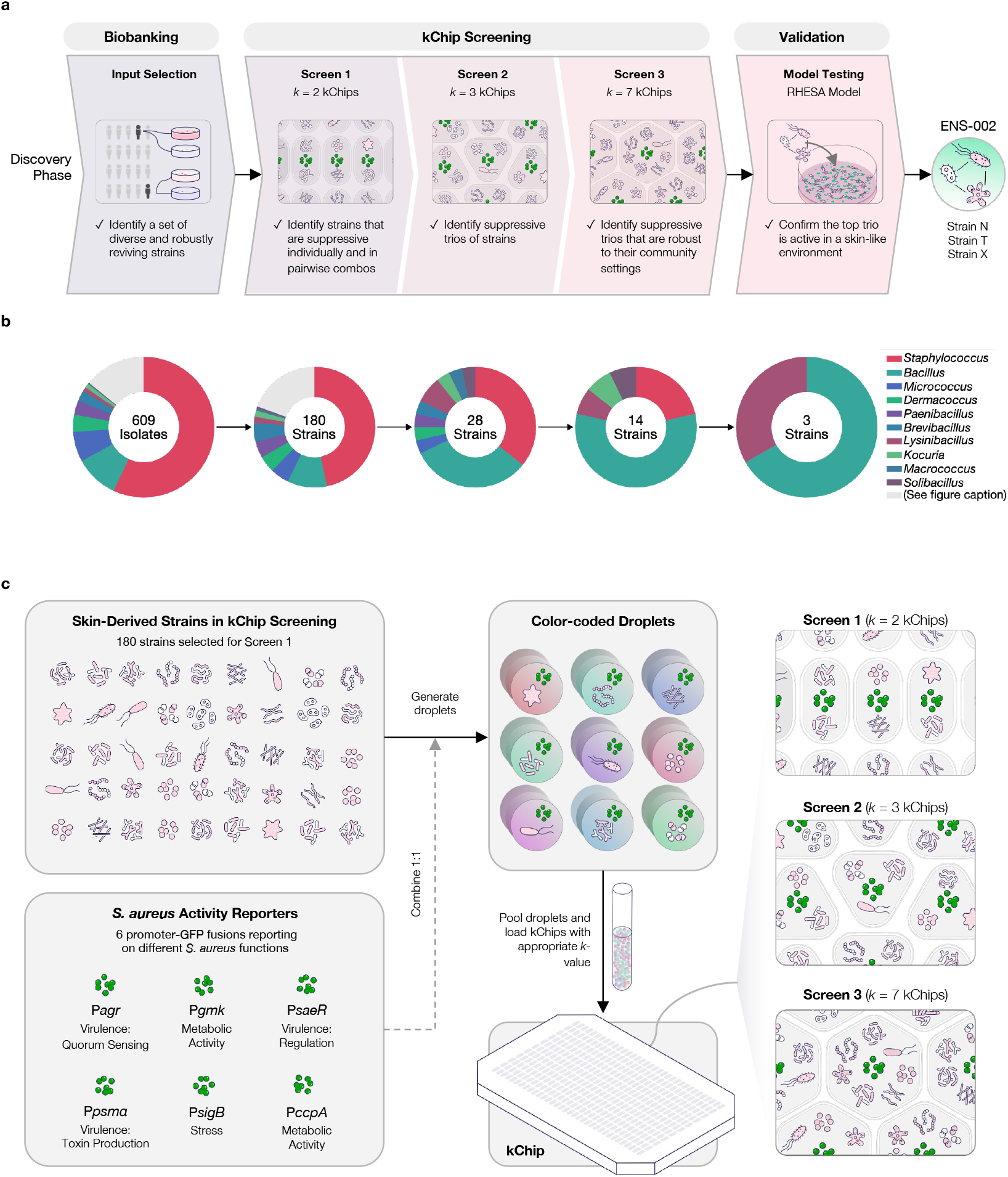
Overview of discovery strategy to identify ENS-002. **a**, The discovery funnel outlining the steps involved in identifying ENS-002, a topical three-strain live biotherapeutic product (LBP), from a biobank of skin bacterial strains. **b**, Pie charts detailing the genus representation of the biobank strains that were used as inputs for each step detailed above in (**a**). Genera that were represented in the top 28 strain selections resulting from Screen 1 are identified by color. For the other 24 genera, see Supplementary Table 1. **c**, An overview of the kChip screening strategy. Cocultures of skin biobank strains and *S. aureus* fluorescent reporter strains were generated, color-coded, dropletized, pooled, and loaded onto a kChip containing microwells that can support two droplets (*k* = 2) for pairwise screening, three droplets (*k* = 3) for three-wise screening, or seven droplets (*k* = 7) for seven-wise screening.

To generate the biobank, skin swabs were first collected from 42 sites (21 healthy donors, two anatomical sites per individual) and streaked on agar. 1,701 colonies were picked and frozen in glycerol (Methods); 609 of these were selected for our biobank based on their ability to consistently revive from frozen stocks in liquid media (a prerequisite for kChip screening and manufacturability). The 609 isolates displayed high phylogenetic diversity (based on 16S sequences), representing 94 distinct species (within the phyla Bacillota, Pseudomonadota, and Actinomycetota) and were enriched for common culturable skin bacteria (e.g., *Staphylococcus epidermidis, Staphylococcus caprae*, and *Staphylococcus capitis*) (Fig. 1b, Supplementary Table 1).

From these 609 isolates, 180 were selected for strain-purifying passaging and kChip screening (Fig. 1b) based on additional filtering steps that limited redundant species from a single donor, corrected for culture-based sampling biases, maximized taxonomic and metabolic diversity, and enriched for strains that displayed inhibitory activity against *S. aureus* virulence in preliminary experiments (Methods).

The three strains ultimately comprising ENS-002 were subjected to a comprehensive safety evaluation, including genome-based screening for virulence factors, antibiotic sensitivity testing, and a literature review for any documented pathogenicity (Methods). These analyses, in conjunction with the healthy status of the donor individuals, supported use of these strains in a topically applied LBP for AD patients.

### kChip enabled large-scale screening of microbial communities for suppression of *S. aureus* metabolic activity and virulence

kChip is a high-throughput combinatorial screening platform that enables the rapid construction of defined microbial cocultures via dropletized inputs that self-assemble into small combinations (e.g. pairwise, three-wise, seven-wise) and merge within microwells, with activities quantified via optical readouts^40^ (Fig. 1c, Methods). In three successive kChip screens, we measured the activity of *S. aureus* across >4 million cocultures consisting of two, three, or seven additional strains, respectively. To assay *S. aureus* activity, we used six “reporters”—*S. aureus* BAA-1717 strains that each carried a plasmid encoding GFP under the control of a promoter linked to a key *S. aureus* gene^42,43^ (Fig. 1c). The GFP fluorescence intensity within a kChip microwell quantitatively reported on a given *S. aureus* characteristic like metabolic activity or toxin production in the context of that microwell’s coculture.

The activity of the six reporters provided a broad view of *S. aureus* physiology across various cocultures, with the *agr* reporter (P*agr-gfp*) providing particularly significant insight into the skin microbiome’s role in modulating AD susceptibility. *S. aureus*’ *agr* quorum-sensing system is a global virulence regulator for expression of toxins and proteases that drive itch, skin barrier disruption, and inflammation associated with the emergence and severity of AD lesions^7–13^. Recent work has also highlighted the importance of *agr* in *S. aureus* skin colonization^44^.

To generate cocultures, each kChip input strain was combined with *S. aureus*, dropletized, and loaded onto a kChip (Methods). Each individual microwell could contain distinct (“AB…”) or repeated strain inputs (“AA…”) depending on the random assembly. Antibiotic and medium-only untreated control inputs were also combined with *S. aureus*. After a 24 h incubation at 30°C or 37°C, the GFP signal from each microwell was measured to quantify *S. aureus* activity relative to untreated monoculture controls (Methods).

Three successive kChip screens, along with off-kChip in vitro assays, whole genome sequencing, and additional phylogenetic characterization, enabled iterative downselection of our biobank of 180 skin strains to the three strains comprising ENS-002 (Fig. 1a, b, Methods).

### Pairwise screening of biobank strains revealed combinations with distinct suppressive effects on *S. aureus* metabolic activity, virulence, and stress

Using kChip, we first aimed to select individual strains that suppressed *S. aureus* metabolic activity and virulence, with priority given to strains that robustly produced strong, if not stronger, suppression in combination with other strains. Screen 1 employed *k* = 2 kChips to evaluate the effects of individual strains AA (i.e. one repeating input) and pairs of distinct strains AB on each of four *S. aureus* reporters: one measuring metabolic activity (P*gmk*-*gfp*), two measuring virulence [quorum sensing (P*agr*-*gfp*), toxin production (P*psmα*-*gfp*)], and one measuring stress (P*sigB*-*gfp*). All 180 strains AA and 16,110 pairs AB were generated in Screen 1 across 3.6M microwells, with a median of 33–36 replicates per combination per reporter.

A range of *S. aureus* responses was observed across the reporters, with 47–71% of combinations producing an undetectable or mild (“neutral”) effect on the given reporter, 20–43% producing an up-to-8-fold decrease (“moderate suppression”), and 9–12% exhibiting an >8-fold decrease (“strong suppression”) (Fig. 2a–d) (Methods). The effects of strains within 16S-determined taxonomic bins were often similar, though we observed considerable within-species and within-genus variance (Fig. 2e, Supplementary Fig. 1–3). For example, the *Bacillus* genus contained many strongly suppressive members across most of the tested combinations; by contrast, a single strain of *Brevibacillus* appeared consistently suppressive unlike other members of its genus.

**Fig. 2.**
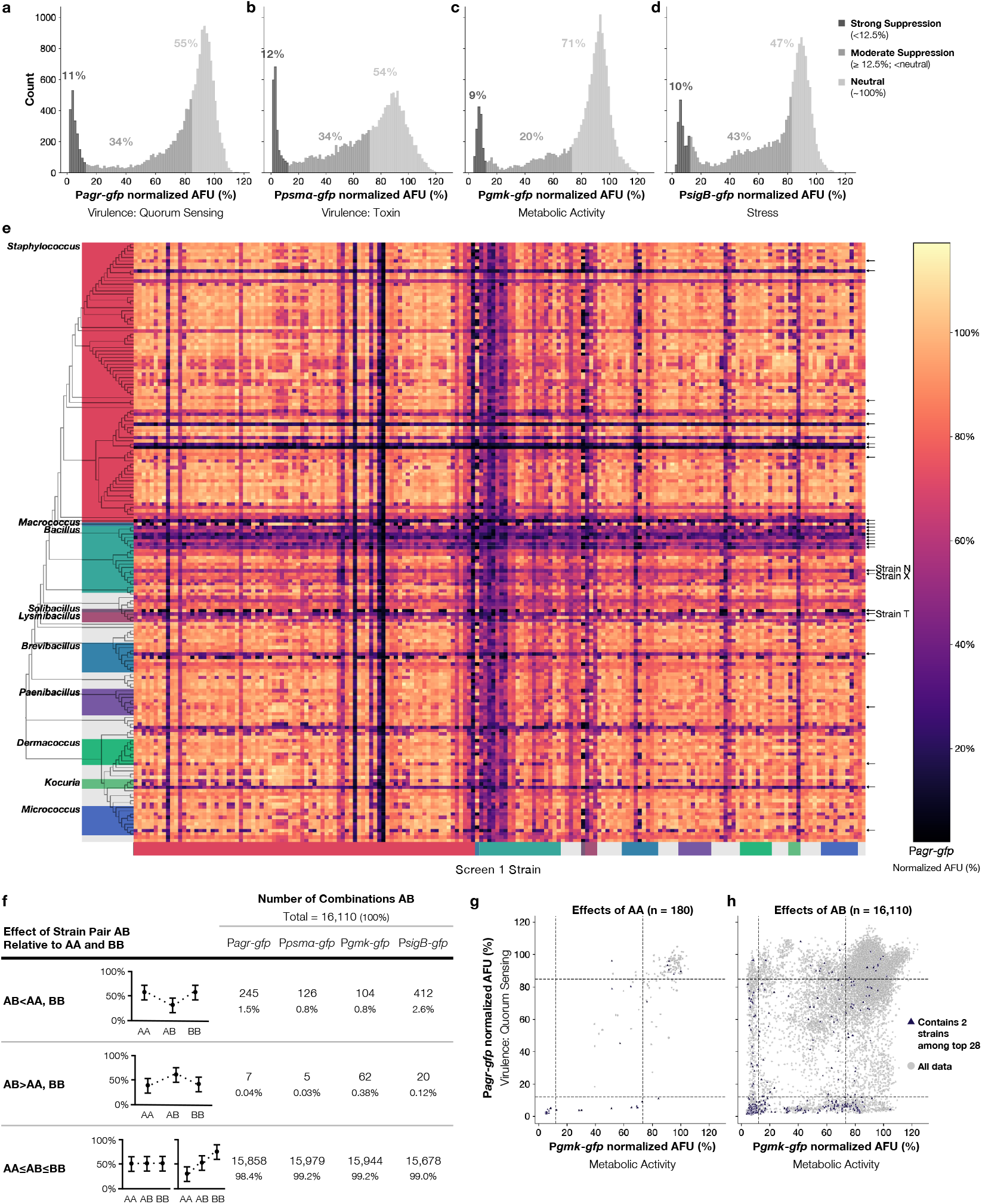
Screen 1: Pairwise screening revealed 28 strains that suppress *S. aureus* alone or in combinations. **a**-**d**, The median effects of each individual strain (AA) and pair or distinct strains (AB) on the measured *S. aureus* reporters P*agr*-*gfp* (**a**), P*psmα*-*gfp* (**b**), P*gmk*-*gfp* (**c**), and P*sigB*-*gfp* (**d**) activity relative to the untreated control after 24 h are shown. The combinations’ effects were categorized as exhibiting “strong suppression” (dark gray, <12.5%), “moderate suppression” (medium gray, ≥12.5%; <neutral), or “neutral” effects (light gray, Methods), with the percentage of pairs in each category indicated above each histogram. **e**, The median relative effect of each pairwise biobank combination on P*agr*-*gfp* activity. A maximum likelihood phylogenetic tree was constructed based on 16S sequences to sort strains. Two of the strains that were screened were later found to be polymicrobial and were not included in the tree. Colored boxes indicate genera of the top 28 strain selections resulting from this screen. Arrows on the right-hand side of the heat map indicate the 28 strains that were selected for further testing; “*” indicates the eventually-selected ENS-002 strains. **f**, The number of combinations AB for each measured reporter that exhibited statistically significantly stronger suppression than the constituent strains (AB<AA, BB) or weaker suppression (AB>AA, BB) or a neutral effect (AA≤AB≤BB) are indicated (based on paired t-tests with a Bonferroni-corrected significance threshold of 1.55 × 10^-6^). **g**-**h**, The median relative effect of each individual biobank strain AA (**g**) or pairwise combination AB (**h**) on P*agr*-*gfp* and P*gmk*-*gfp* activity. Blue triangles indicate the strains or combinations containing two inputs of the 28 strains selected for advancement; gray circles indicate all remaining data. Dashed horizontal and vertical lines indicate bounds for strong and moderate suppression. Data shown are median effects from a median replicate count of 35 (P*agr*), 32 (P*psmα*), 33 (P*gmk*), and 36 (P*sigB*). AFU, arbitrary fluorescence units.

The degree of *S. aureus* suppression by a given strain pair AB was usually within the range of the suppressive effects of AA and BB (98.4–99.2% across the four reporters) with notable exceptions (Fig. 2f). In 0.8–2.6% of combinations, we observed stronger suppression than either constituent (Fig. 2f). Rarer still were the 0.03–0.38% of combinations that were less suppressive than either of their constituents.

Strong suppression of more than one *S. aureus* reporter was common. Of the four reporters tested, twenty individual strains strongly suppressed (>8-fold decrease) at least one reporter, 17 strains at least two reporters, and eight strains all four reporters. Cross-reporter relationships are depicted in Fig. 2g,h and Supplementary Fig. 4. Notably, strong P*agr* suppression and neutral P*gmk* activity were observed with only one AA strain but 462 AB pairs (Fig 2g,h). Of these 462, 163 (35%) did not contain any of the 18 strains that individually strongly suppressed P*agr*.

A total of 28 strains were selected for advancement. The 17 strains that individually exhibited strong suppression of two or more *S. aureus* reporters were all selected. While optimizing for overall genetic diversity, 11 additional strains that were prevalent in pairs more suppressive than their constituents (typically of three or more *S. aureus* reporters) were selected. These 28 strains represented nine genera, including 10 *Staphylococcus* and 9 *Bacillus* strains (Fig. 1b). AA strains and AB pairs containing these 28 selections are highlighted in Fig. 2e,g,h.

### Three-wise combination screening enabled selection of 14 top-performing *S. aureus*-suppressing strains

To identify suppressive trios among the 28 Screen 1 selections, Screen 2 measured the effects of three-wise combinations in *k* = 3 microwells on each of six *S. aureus* reporters strains: two measuring metabolic activity (P*gmk-gfp*, P*ccpA-gfp*), three measuring virulence [quorum sensing (P*agr-gfp*), toxin production (P*psmα-gfp*), and regulation (P*saeR-gfp*)], and one measuring stress (P*sigB-gfp*). The screen generated data from microwells containing randomly assembled three-wise combinations, covering all 4,060 possible strain combinations. Inputs could repeat with any multiplicity (e.g., AAA, ABB, ABC); we focused on the 3,276 non-repeating combinations ABC and single strains A[][] (where [][] indicates medium-only control inputs, as instances of AAA, or one repeating input, were statistically rarer given random assembly), which were measured from >700,000 microwells with a median of 31–40 replicates per combination per reporter (Fig. 3a).

**Fig. 3.**
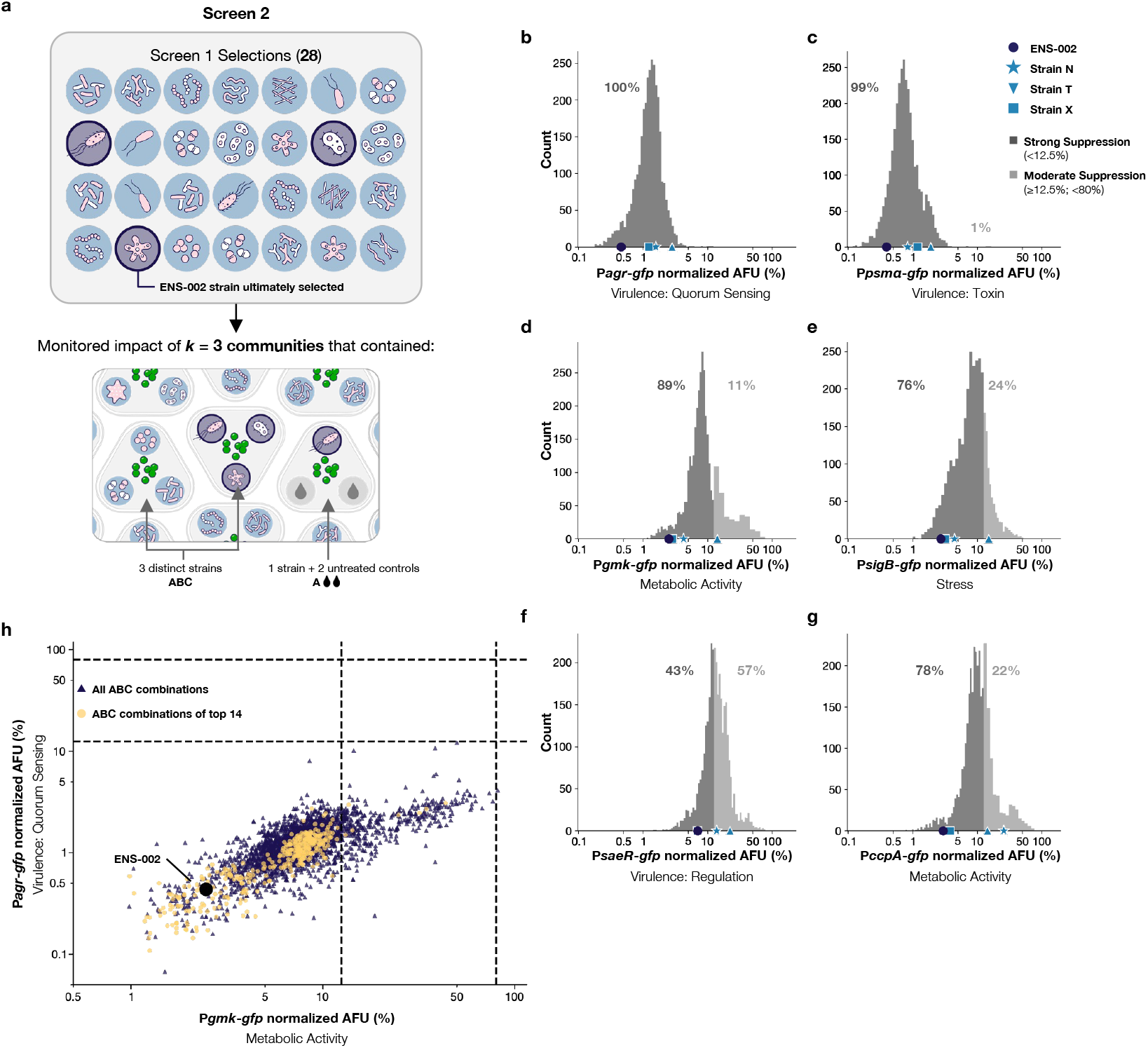
Screen 2: Fourteen top-performing strains emerged from three-wise kChip screening. **a**, A schematic depicting the types of three-wise combinations of interest in (**b**-**g**) that were generated from the 28 Screen 1 selections in Screen 2. Dark blue highlighting indicates the final three ENS-002 strains. Relative *S. aureus* reporter activity with the following types of combinations was monitored: all possible combinations of three distinct Screen 1 selections (ABC, left), including those that received the three strains comprising ENS-002 (middle); and combinations containing one Screen 1 selection with two inputs of untreated controls (A[][], right). Other combinations that were assembled were excluded from analyses. **b**-**g**, The median effects of all ABC combinations on the measured *S. aureus* reporters P*agr*-*gfp* (**b**), P*psmα*-*gfp* (**c**), P*gmk*-*gfp* (**d**), P*sigB*-*gfp* (**e**), P*saeR*-*gfp* (**f**), and P*ccpA*-*gfp* (**g**) activity relative to the untreated control after 24 h are shown. The effects were categorized as exhibiting “strong suppression” (dark gray, <12.5%) or “moderate suppression” (medium gray, ≥12.5%; <80%) with the percentage of combinations in each category indicated above each histogram. The symbols along the X-axes indicate the median effect of microwells receiving all three ENS-002 strains (circles) or one input of each strain with 2 medium-only controls (A[][], Strain N: star; Strain T: triangle; Strain X: square). Suppression by ENS-002 was significantly stronger than by Strain T for all tested reporters; stronger than by Strain N for P*agr*, P*psmα*, and P*ccpA*; and stronger than Strain X for P*agr* and P*psmα* (*P* <0.017, two-sided Mann-Whitney U test with Bonferroni correction). **h**, The median relative effect of all non-repeating three-wise combinations of biobank strains ABC on P*agr*-*gfp* and P*gmk*-*gfp* activity. Yellow circles indicate the combinations that received three non-repeating inputs of the 14 strains selected for advancement; blue triangles indicate all remaining data. The ENS-002 combination is indicated with the black circle. Dashed horizontal and vertical lines indicate bounds for strong and moderate suppression depicted in **b**-**g**. See Supplementary Fig. 5 for the remaining cross-reporter scatter plots. Data shown are median effects from a median replicate count of 34 (P*agr*), 37 (P*sigB*), 31 (P*psmα*), 35 (P*saeR*), 40 (P*ccpA*), and 35 (P*gmk*). AFU, arbitrary fluorescence units.

Of the 3,276 ABC combinations, 76–100% strongly suppressed (>8-fold decrease) the P*agr*, P*psmα*, P*gmk*, P*sigB*, and P*ccpA* reporters (Fig. 3b–g), which was far more frequent than in Screen 1. (Only 43% of combinations strongly suppressed the P*saeR* reporter, consistent with the fact that P*saeR* suppression was not included as a selection criterion in Screen 1.)

From Screen 2 data, 14 top-performing strains were selected for advancement for their appearance in combinations that strongly suppressed *S. aureus*. Strong suppression against multiple reporters was prioritized (with some exclusions made due to preliminary safety assessments based on taxonomic assignments) (Fig. 1b, Methods). In selecting 14 of the 28 strains, the combinatorial space was reduced to 11% of possible non-repeating trios (from 3,276 to 364). Among these 364, 357 (98%) strongly suppressed both the *S. aureus* P*agr* and P*gmk* reporters (Fig. 3h). A similar suppression pattern was observed with the other reporter combinations (Supplementary Fig. 5).

While its composition was later determined, the Screen 2 performance of ENS-002 is highlighted in Fig. 3b–h. Among the 364 trios, ENS-002 ranked in the 88th percentile or higher for suppression of each reporter. Suppression of each measured reporter by ENS-002 was as strong or stronger than that of its constituents. ENS-002 suppressed two *S. aureus* virulence reporters [quorum sensing (P*agr*) and toxin production (P*psmα*)] significantly more strongly (*P* <0.017, Methods) than any of its constituent strains alone; suppression of P*gmk* and P*saeR* by ENS-002 and Strain X was not statistically distinct.

### The three-strain consortium ENS-002 strongly suppressed *S. aureus* across varied bacterial communities

To determine how robustly the 364 trios among the 14 Screen 2 top-performing strains suppressed *S. aureus* when embedded within larger microbial communities, Screen 3 measured seven-wise combinations on *k* = 7 kChips. Strains were drawn from a set of 92: the 14 Screen 2 selections and 78 diverse skin community member strains (that were excluded from, or did not advance past, Screen 1) (Fig. 4a, Supplementary Fig. 6). For each community, we measured *S. aureus* virulence [quorum sensing (P*agr*-*gfp*) and toxin production (P*psmα*-*gfp*)] or metabolic activity (P*gmk*-*gfp*). We focused our analyses on microwells that received exclusively community members (MMMMMMM, which occurred in 1,618–1,990 microwells for each reporter and allowed repeating members); and those that received exactly three non-repeating Screen 2 selections and four community members (ABCMMMM, which occurred in 2,222–3,282 microwells for each reporter) (Fig. 4a).

**Fig. 4.**
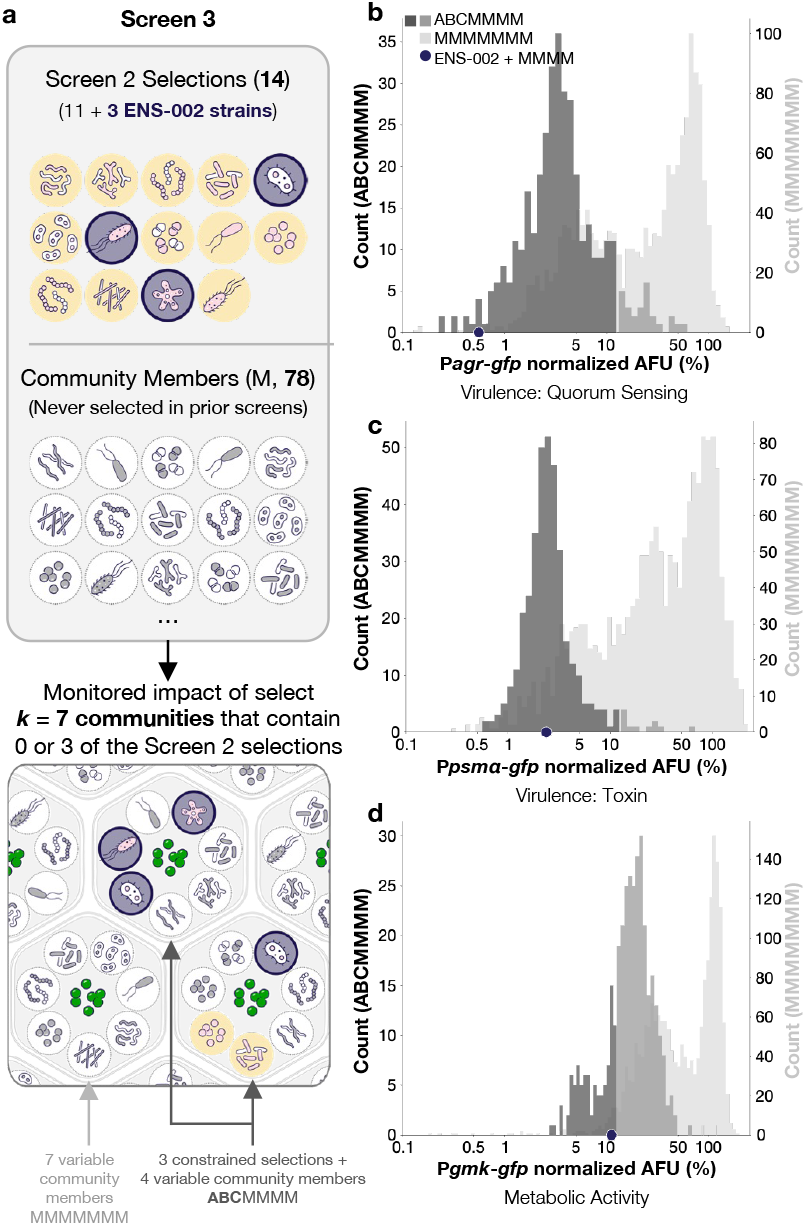
Screen 3: ENS-002 maintained strong *S. aureus*-suppressive activity within the context of varied bacterial communities. **a**, A schematic depicting the types of seven-wise combinations of interest in (**b-d**) that were generated from the 14 strain selections from Screen 2 (yellow highlighting; dark blue highlighting for ENS-002) and 78 community members (see Supplementary Fig. 6) (gray dashed circles). **b**-**d**, For the *S. aureus* reporters P*agr*-*gfp* (**b**), P*psmα*-*gfp* (**c**), and P*gmk*-*gfp* (**d**), two distributions of *S. aureus* activity relative to the untreated control are shown. The background light gray distribution shows the relative *S. aureus* activity of all assembled MMMMMMM combinations (counts on right axes). In the case a particular community was assembled more than once, the median is reported. The forefront distribution shows the median relative *S. aureus* activity of all assembled ABCMMMM combinations (counts on left axes) [dark gray for strong suppression (<12.5%); medium gray for moderate suppression (≥12.5%)]. For the P*agr*, P*psmα*, and P*gmk* reporters, data were collected from 1618, 1760, 1990 MMMMMMM microwells and 2222, 3282, and 2792 ABCMMMM microwells, respectively. The blue circles along the X-axes indicate the median reporter activity observed in microwells receiving all three ENS-002 strains and four community members (data collected from 4, 7, and 5 microwells for the P*agr*, P*psmα*, and P*gmk* reporters, respectively). AFU, arbitrary fluorescence units.

ABCMMMM combinations exhibited higher levels of *S. aureus* suppression (2–17% of the untreated controls) than MMMMMMM combinations (27–59%) (Fig. 4b–d). Of the 364 strongly suppressive (>8-fold decrease) Screen 2 combinations ABC, 96% remained strongly suppressive of P*agr*, 100% of P*psmα*, and 26% of P*gmk* across variable communities ABCMMMM.

The trio ENS-002 emerged as a top combination (Fig. 4b–d). Communities containing ENS-002 suppressed the P*agr*, P*psmα*, and P*gmk* reporters to 0.6%, 2.4%, and 11.2% (98th, 51st, and 80th percentile), respectively.

ENS-002 constituents Strain N, Strain T, and Strain X each individually suppressed P*agr* to <4% (85th percentile and above), P*psmα* to <7% (39th percentile and above), and P*gmk* to <25% (69th percentile and above) (Supplementary Fig. 7).

### ENS-002 suppressed *S. aureus* virulence, metabolic activity, and viability in microtiter plate assays

ENS-002-mediated suppression of *S. aureus* virulence (P*agr*-*gfp*) and metabolic activity (P*gmk*-*gfp*) observed on kChip was validated through microtiter plate liquid coculture assays. ENS-002-treated cultures exhibited marked suppression; all three individual strains and the combination decreased expression from the P*agr* reporter to approximately the lower limit of detection (Fig. 5a). Effects on P*gmk* were more variable: Strain N, Strain X, and the ENS-002 combination reduced expression to ∼4–6% of the untreated PBS control, while Strain T reduced it to 15% (Fig. 5b). Treatment with wild type *S. aureus* decreased expression by both reporters to roughly half that of the untreated control.

**Fig. 5.**
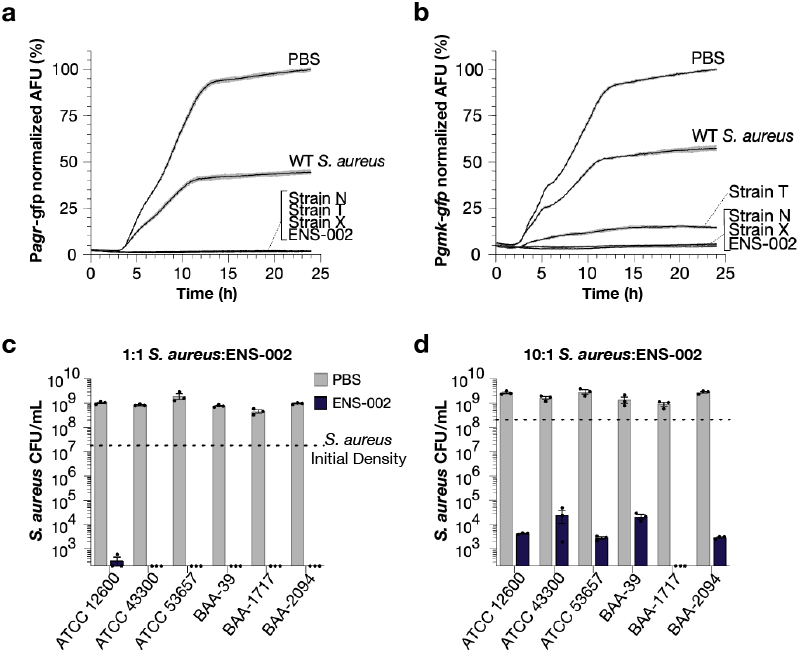
ENS-002 decreased *S. aureus* virulence, metabolic activity, and load in microtiter plate assays. **a-b**, Cocultures of the *S. aureus* strain with the P*agr*-*gfp* (**a**) or P*gmk*-*gfp* (**b**) reporters with either: ENS-002, its individual constituent strains (Strain N, Strain T, or Strain X), or the non-fluorescent parent *S. aureus* strain in a 1:1 CFU ratio, or a PBS control were generated in microtiter plates. GFP fluorescence was measured every 7.5 min for 24 h. GFP measurements were normalized to the median endpoint untreated GFP value and are connected by the solid black lines; SEM is indicated by gray shading above and below. **c**,**d**, Six *S. aureus* clinical isolates, transformed with the plasmid encoding the P*gmk*-*gfp* reporter construct, were cocultured with ENS-002 in either a 1:1 (**c**) or 10:1 (**d**) *S. aureus*:ENS-002 CFU ratio or a PBS control. *S. aureus* CFU/mL after 24 h are reported. Bars indicate mean ± SEM of three replicates; light gray is untreated, dark gray is treated with ENS-002. The mean *S. aureus* starting density at T = 0 h is indicated by the dashed horizontal lines (1.9 ⍰ 10^7^ CFU/mL and 2.1 1 10^8^ CFU/mL, respectively). BAA-1717 is the same parent strain used for all other reported data. AFU, arbitrary fluorescence units; CFU, colony forming units.

We further demonstrated that ENS-002 impacts viable *S. aureus* load across multiple *S. aureus* strains via a microtiter plate colony forming unit (CFU) enumeration assay. We tested how ENS-002 affected the viability of *S. aureus* BAA-1717 (kChip screening strain) and, because different *S. aureus* strains dominate across AD patients^45^, five additional *S. aureus* strains isolated from human patients (Methods). For a 1:1 *S. aureus*:ENS-002 ratio test, the *S. aureus* density dropped to near or below the limit of detection for all tested *S. aureus* strains. This drop constituted a >10^5^-fold CFU decrease relative to untreated controls and a >10^4^-fold decrease relative to the starting inoculum (Fig. 5C). Even at a 10:1 *S. aureus*:ENS-002 ratio, the *S. aureus* density decreased >10^4^-fold relative to the untreated control and starting inoculum (Fig. 5D).

### ENS-002 exhibited anti-*S. aureus* activity on a reconstructed human epidermis (RHE) model

To evaluate ENS-002 in an environment that more closely mimics human skin, we developed the RHESA (Reconstructed Human Epidermis *S. aureus* Activity) model. We pre-seeded the P*agr*-*gfp S. aureus* reporter strain onto EpiDerm™ tissue models (MatTek, Ashland, MA) and monitored the effects of applied treatments on *S. aureus* CFU and P*agr*-*gfp* after 24 h (Fig. 6a; see Methods and Supplementary Fig. 8).

**Fig. 6.**
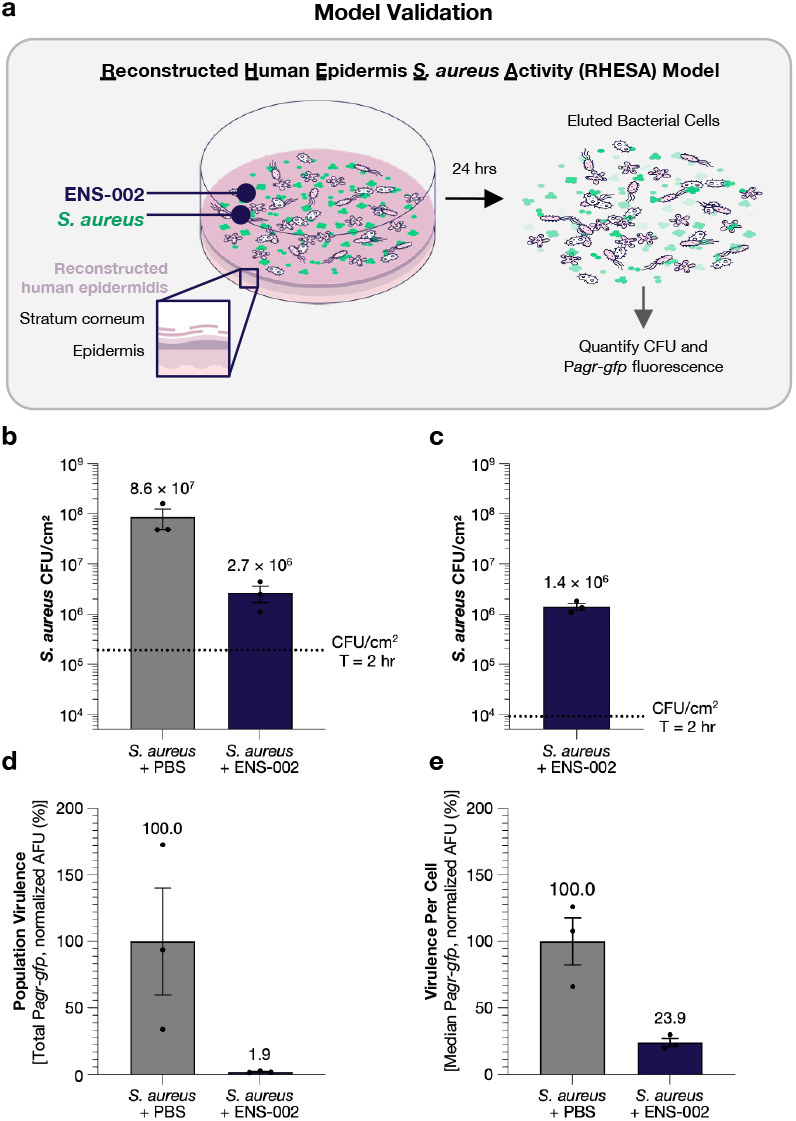
ENS-002 decreased the *S. aureus* load and virulence on reconstructed human epidermis (RHE) models. **a**, A cartoon representation of the RHESA (Reconstructed Human Epidermis *S. aureus* Activity) model workflow. RHE models (MatTek, EPI-100-NMM-HCF-AFAB) are pre-seeded with the *S. aureus* P*agr*-*gfp* reporter strain and then treated with ENS-002. After a 24 h incubation at 37°C, 5% CO_2_, bacterial cells are eluted, plated for enumeration, and analyzed by flow cytometry to quantify virulence of the *S. aureus* cells. See Supplementary Fig. 8 for a more detailed timeline. **b**,**c**, *S. aureus* (**b**) and ENS-002 (**c**) enumeration at T = 24 h. The dashed horizontal lines indicate the *S. aureus* and ENS-002 densities (1.9 ⍰ 10^5^ CFU/cm^2^ and 9.1 ⍰ 10^3^ CFU/cm^2^, respectively) at T = 2 h (the earliest measurable time point, Methods). **d**,**e**, The population-level (total) GFP fluorescence (GFP) (**d**) and median GFP fluorescence (GFP) per fluorescent cell (**e**) normalized to the average of the PBS-treated controls, detected by running a set volume of the eluted samples through an Attune CytPix Flow Cytometer at T = 24 h. See Supplementary Fig. 9 for fluorescence distributions. Bars indicate mean ± SEM of three replicates, with the mean labeled above each bar. AFU, arbitrary fluorescence units; CFU, colony forming units.

We tested ENS-002 activity on the RHESA model using a ∼20:1 inoculation ratio of *S. aureus* to ENS-002 (2.0 × 10^5^ CFU/cm^2^ and 9.1 × 10^3^ CFU/cm^2^, respectively). After the 24 h incubation, *S. aureus* on untreated models underwent 8.8 doublings (yielding 8.6 × 10^7^ CFU/cm^2^ on average across three replicates) (Fig. 6b). In contrast, *S. aureus* on models treated with ENS-002 underwent 3.8 doublings (yielding 2.7 × 10^6^ CFU/cm^2^ on average across three replicates), yielding 30-fold fewer CFU than observed with the untreated controls. Collectively, ENS-002 cells underwent 7.3 doublings (yielding 1.4 × 10^6^ CFU/cm^2^ on average across three replicates) (Fig. 6c).

ENS-002 treatment also corresponded with a decrease in the measured virulence of the remaining *S. aureus* population both at the population and individual cell level. Measured via flow cytometry, the population-level virulence of the *S. aureus* cells treated with ENS-002 decreased >50-fold (to 1.9% of the untreated control) (Fig. 6D). The average virulence per fluorescent cell that remained decreased >4-fold (to 23.9% that of the untreated control) (Fig. 6E, Supplementary Fig. 9).

## Discussion

*S. aureus* plays a significant role in the pathogenesis of AD, exploiting the compromised skin barrier, inducing inflammation, and perpetuating immune dysregulation. Existing AD therapies do not directly address *S. aureus*’ contributions to the disease. We present the discovery of ENS-002, a combination of three bacterial strains sourced from healthy human skin that robustly suppresses *S. aureus* virulence and proliferation, both of which are etiologically implicated in AD flares. The strong performances of ENS-002 strains individually and in combination throughout the described kChip screens, microtiter assays, and RHESA model experiments, combined with safety and manufacturability evaluations (Methods) ultimately led to their selection for further development as a therapeutic topical LBP for treating AD.

Our combinatorial datasets revealed that strain combinations drive a wider variety of specific *S. aureus* physiological responses than individual strains. For instance, Screen 1 revealed pairwise combinations with distinct effects from individual strains on the tested *S. aureus* reporters, including those that strongly suppressed P*agr* but neutrally affected P*gmk* (suggesting a specific growth-independent suppression of quorum sensing) (Fig. 2g,h). The screening throughput required to identify these distinct combinatorial outcomes was on the order of millions of data points.

Screening at this scale also facilitated the identification of combinations like ENS-002 that exhibited stronger suppression than that of their constituents (Fig. 2f and Fig. 3b,c). In these cases, it’s possible that suppressive mechanisms acted on *S. aureus* without interference or with synergy. The differential responses among the *S. aureus* reporters also suggested distinct mechanisms of suppression among members of strongly suppressive combinations. For example, unlike Strains N and X, Strain T strongly suppressed P*agr* without full suppression of P*gmk* (Fig. 5a,b). Additional work is required to discern the molecular basis for these observed effects on *S. aureus*.

In principle, ENS-002 can benefit more patients if it can suppress *S. aureus* in a manner that is robust to microbiome diversity within and among hosts. When tested across variable microbial community compositions on kChip, ENS-002 robustly suppressed both virulence and metabolic activity (Fig. 4b–d). Robust inhibition of metabolic activity was a critical distinguishing quality of ENS-002: Among the 14 top isolates, 98% of trios strongly suppressed metabolic activity (Screen 2) but only 26% retained this ability in larger communities (Screen 3). ENS-002’s robustness presages successful activity across diverse skin microbiomes. We hypothesize that the inclusion of three strains in an LBP increases the chances that one or more of the constituent strains will retain suppressive activity in different contexts, which may also bolster robustness.

ENS-002 strains, by virtue of being isolated from human skin, may be well-adapted to the skin milieu, enabling long-term LBP activity. Indeed, ENS-002 suppressed *S. aureus* on our RHESA tissue culture model, a skin-like, nutrient-limited environment (Fig. 6b–e). As the starting *S. aureus* density loaded onto the models fell into the 90th percentile of that observed in AD lesional skin^32^, we expect that ENS-002 can suppress *S. aureus* at levels relevant to the skin of AD patients.

ENS-002’s therapeutic potential extends to *S. aureus* decolonization. It’s been shown that disrupting *S. aureus* Agr quorum sensing, which appears critical for initiating and maintaining *S. aureus* colonization^44,46^, can be an effective strategy for decolonization^46^. The observed >50-fold population-level P*agr* suppression on the RHESA model with ENS-002 treatment (Fig. 6d,e), combined with the >10^4^-fold *S. aureus* viability decrease in ENS-002 cocultures (Fig 5c,d), suggests that ENS-002 may be able to decolonize *S. aureus* altogether. As the benefits of *S. aureus* decolonization extend beyond AD treatment, ENS-002 may be an effective LBP for multiple *S. aureus*-implicated skin conditions like hand eczema, impetigo, folliculitis, and skin barrier dysfunction^47–49^.

While kChip has previously been used to screen interactions among synthetic microbial communities^40,50^, this is the first reported example of the use of kChip in the discovery of an LBP. Accordingly, this is also the first example of an LBP discovery process that relied on the generation of millions of synthetic microbial communities to select the most optimal combination from a set of hundreds of strains. The screening strategy outlined here has the potential to be used for the discovery of microbial products that address the microbial drivers of other skin conditions like body odor, acne, and dandruff^51–53^.

The primary goal of this work was to discover a consortium of bacteria that strongly suppresses *S. aureus* growth and virulence to treat AD. Via massive coculture screening, a consortium was discovered consisting of bacterial species that, to our knowledge, have not yet been investigated for AD treatment. ENS-002 was therefore further developed and tested in a Phase I study, “Topical ENS-002 for Atopic Dermatitis in Adults (EnSync)” (NCT06469385), to determine its safety and effects, and is now undergoing development for a Phase II study.

## Methods

### Skin bacterial biobank assembly

A skin bacterial biobank was generated by swabbing the skin of healthy volunteers. Twenty-one volunteers were selected as skin swab donors for biobank generation after self-reporting any history of skin disease, autoimmune disease, and drug use; swabs collected from anyone with systemic or topical antibiotic use in the prior six months were excluded from the biobank. Volunteers were asked to avoid showering 24 hours prior to the sample collection. Two sites per volunteer (volar forearm and antecubital fossa—predominant sites of AD) were swabbed. Resuspensions of these swabs were grown on 7–12 agar-based media; resulting individual colonies were selected as isolates for the biobank. This protocol was reviewed and approved by WCG IRB (IRB Tracking Number 20201725). Informed consent was obtained from all participants.

In total, 1,701 skin isolates were selected and saved in glycerol stocks. Of these, 609 isolates were selected for further evaluation based on their ability to be revived from glycerol stocks (a preliminary manufacturability assessment) in TSB while simultaneously attempting to deduplicate strains based on colony morphology. These isolates were submitted for 16S amplification and Sanger sequencing (Genewiz/Azenta Life Sciences). Taxonomic assignments were identified via NCBI’s BLAST (Basic Local Alignment Search Tool) in February 2021; 43 failed sequencing and 15 were of poor quality (% identity on the top BLAST hits were <97% and reads <750 nt) and were later re-submitted for sequencing and identified in September 2025; five of these isolates were found to contain multiple species and were accordingly classified as “mixed”. These taxonomic assignments were used for preliminary safety assessments during Screen 2 top-performing strain selection; putative pathogens (based upon taxonomic assignment) were generally excluded from advancement.

A diverse subset of isolates was selected for kChip screening based on taxonomic assignments and preliminary assessments of their capabilities. The metabolic capabilities of these isolates were evaluated by measuring the impact of a panel of carbon sources at 0.5% (w/v) on outgrowth in glucose-free tryptic soy broth (TSB BD 286220), used at a 25% (v/v) concentration relative to the manufacturer’s instructions. The effects of these isolates on four *S. aureus* reporters (P*agr*, P*psmα*, P*sigB*, and P*gyr*) in microtiter plates was also measured. Based on taxonomic assignments, the ability to grow with diverse carbon sources, and preliminary observed *S. aureus* suppression, 180 isolates were selected for kChip screens. The 180 biobank isolates used for kChip screening underwent one strain-purifying passage, in which a single colony was selected from a TSA plate, before being used for screening. Two isolates were not successfully strain-purified at this stage and consequently still received a taxonomic assignment of “mixed”. Strains N, T, and X underwent two additional strain-purifying passages prior to post-kChip screening evaluations. The commensal community strains used for Screen 3 were not passaged before use.

### Whole genome sequencing, safety, and manufacturability analyses of ENS-002 strains

To support the expectation that Strains N, T, and X, which were isolated from healthy human skin, should be safe for application to patient skin, their genomes were sequenced and analyzed, their antibiotic susceptibilities were assessed, and a literature search was performed based on their taxonomic assignments.

Whole genome sequencing and assembly was performed by CosmosID (Germantown, MD). Taxonomic assignments were made using the MUMmer alignment database^54^. Targeted searches using the VFDB in November 2023^55,56^ did not detect virulence factors in these whole genome sequences.

Minimum inhibitory concentration (MIC) testing, which was performed at BA Sciences (Salem, NH) according to guidance from the Clinical Laboratory Standards Institute (CLSI), identified several overlapping antibiotic susceptibilities of the three strains. This outcome indicates that, if necessary, all three strains in ENS-002 could be killed with a single or combination antibiotic regimen.

A literature search was performed using Google Search in March 2024, combining the species names of Strains N, T, and X, or closely related species, with the terms: 1) “Infection,” 2) “Disease,” 3) “Pathogenicity,” 4) “Virulence,” 5) “Clinical,” 6) “Case study,” 7) “Safety,” or 8) “Human.” The literature searches did not reveal any reported instances of infection or disease caused by these species.

A manufacturability assessment was also performed to confirm that growth of the ENS-002 strains was scalable and that *S. aureus*-suppressive activities were maintained when grown in animal-derived component free (ACDF) media for subsequent cGMP production (data not shown).

### *S. aureus* reporter generation

Six plasmid-encoded reporters were constructed to produce GFP under the control of promoters regulating key *S. aureus* genes related to virulence [quorum sensing (P*agr-gfp*), toxin production (P*psmα-gfp*), and regulation (P*saeR-gfp*)]^13^, metabolic activity [housekeeping (P*gmk*-*gfp*) and carbon catabolism regulation (P*ccpA*-*gfp*)]^43,57^, and stress (P*sigB*-*gfp*)^58^. First, an entry vector was constructed by combining the PCR-amplified rep and chloramphenicol resistance genes from pC194 (isolated from ATCC 37034, a plasmid that can replicate in *S. aur*eu*s*^57^) with the synthesized genes encoding sfGFP and chloramphenicol resistance based on pYTK001^60^ (for intermediate cloning in *E. coli*) via Gibson assembly^61^. Then, PCR amplicons (from *S. aureus* ATCC 35556) of the desired promoter regions were inserted into the entry vector using Golden Gate assembly with BsmBI and T4 DNA Ligase^62^. The boundaries of promoters are summarized in Supplementary Table 2. The plasmids were separately transformed into *S. aureus* strain ATCC BAA-1717, a USA300 strain isolated from an adolescent patient, following previously described protocols^63^. Unless otherwise indicated, the *S. aureus* reporter strains used in each experiment were those constructed in the BAA-1717 background. The data shown in Fig. 5c–d are the exception and were generated from five other *S. aureus* strains (ATCC 12600, ATCC 43300, ATCC 53657, BAA-39, and BAA-2094, all isolated from human patients) transformed with the plasmid containing the P*gmk*-*gfp* reporter.

### Bacterial culturing

All strains underwent a pre-growth period prior to the start of an experiment: strains were grown on tryptic soy agar plates (TSA; BD 236950) and/or in glucose-free TSB. *S. aureus* cultures were grown with chloramphenicol (10 μg/mL) to ensure reporter plasmid maintenance. For kChip screens, glycerol stocks of strains were inoculated directly into the glucose-free TSB. For off-kChip experiments, strains were first streaked from glycerol stocks onto TSA plates; single colonies were used to inoculate liquid cultures.

Prior to the start of an experiment, *S. aureus* cultures were grown at 30°C (for Screens 1 and 2) or 37°C (for Screen 3 and all other work) for 24 h with shaking (250 RPM); biobank strains were grown at 30°C (for Screens 1 and 2) or 37°C (for Screen 3 and all other work) for 24 h without shaking for kChip screening; for all off-kChip experiments, Strains N, T, and X were grown at 37°C for 24 h with shaking (250 RPM); for RHESA model experiments, cultures of Strains N, T, and X cultures were back-diluted 1:100 into glucose-free TSB and grown with shaking (250 RPM) at 37°C for 3.5 hr, such that they would be loaded in exponential-phase onto the RHESA models. Cultures were all normalized by optical density at 600 nm (OD_600_) to the desired starting density for each experiment.

CFU were measured post-normalization and at the desired endpoints for off-kChip coculture and RHESA model experiments. Each culture was serially diluted and spot-plated onto TSA plates to quantify ENS-002 CFU and onto TSA plates containing 10 μg/mL chloramphenicol to quantify *S. aureus* CFU. TSA plates without antibiotics were incubated at RT for 24 h before enumeration; TSA plates containing chloramphenicol were incubated at 30°C for 24 h before enumeration. To quantify ENS-002 on TSA plates, fluorescent *S. aureus* colonies were not included.

### kChip manufacturing

“*k* = 2”, “*k* = 3”, and “*k* = 7” kChips, containing approximately 120,000 2-droplet microwells, 86,000 3-droplet microwells, or 42,000 7-droplet microwells, respectively, were used for these experiments (Fig. 1c). Custom silicone wafer molds were created by photolithography (Custom Nanotech). The kChips were fabricated by soft lithography from polydimethylsiloxane (PDMS; Dow Corning Sylgard) and coated with 1.5 μm parylene C (Kayaku Advanced Materials, formerly Paratronix).

### kChip input preprocessing

Pre-growth cultures of *S. aureus* reporter strains and the biobank strains were diluted before use in the kChip screens: *S. aureus* reporters were normalized to achieve a final OD_600_ = 0.02 in 200 μL for all screens (roughly corresponding to 4 × 10^7^ CFU/mL or an equivalent of 40 CFU in each ∼1 nL droplet); biobank strains were diluted 1:100 (v/v) for Screen 1 and normalized to achieve a final OD_600_ = 0.02 in 200 μL for Screens 2 and 3. kChip input cocultures were assembled in glucose-free TSB and consisted of an *S. aureus* reporter strain culture combined with an individual skin strain culture, a streptomycin antibiotic control, or a medium-only untreated control. The streptomycin concentration was adjusted by the given screen’s *k* value to achieve a final concentration of 100 μg/mL streptomycin if a microwell received one droplet containing the antibiotic control. Every input received a “color code,” or unique ratio of three AlexaFluor fluorescent dyes (AF555, AF594, and AF647 standardized to a final total dye concentration of 2.5 μM) before generating droplets to be used as inputs.

### Droplet making and kChip loading

A similar protocol for preparing droplets, loading, and imaging the kChip to that previously described^40^ was followed. Droplets (1 nL) were produced on a Bio-Rad QX200 Droplet Generator in a fluorocarbon oil (3M Novec 7500). Droplets were pooled to prepare a droplet suspension and injected into a custom-built kChip loading apparatus. Each microwell randomly sampled 2, 3, or 7 droplets. The kChips for Screens 1 2 were incubated at 30°C for 24 h between imaging timepoints on the microscope; the kChips for Screen 3 were incubated at 37°C for 24 h.

### Fluorescence microscopy imaging

All fluorescence microscopy was performed using a Nikon Ti-E inverted fluorescence microscope with fluorescence excitation by a SOLA SE II 365 Light Engine. Images were collected by means of a Hamamatsu ORCA-Flash 4.0 CMOS camera. Images were taken across four fluorescence channels—three for the color codes and one additional channel for the GFP assay with appropriate sets of excitation and emission filters. The signals corresponding to each dye channel were used to identify the contents of a given droplet within each microwell prior to droplet merging (“pre-merge”). The GFP channel was used to quantify the assay signal. The kChip was scanned at 2× magnification to identify the droplets in each microwell from their color codes. Each kChip was imaged in 198 tiles that would be analyzed during data analysis. Droplets were merged within their microwells via ∼10 s of exposure to an alternating-current electric field (4.5 MHz, 10,000–45,000 volts; Electro-Technic Products corona treater)^40^. After a 24 h incubation, the post-merge kChip was scanned at 2× magnification to quantify the GFP signal in each microwell.

### kChip image analysis

A custom image analysis pipeline was used to: 1) Identify droplets within the images; 2) Decode the contents of each droplet based on the color code; 3) Assign each droplet to a microwell; 4) Filter out microwells that did not meet quality control standards to ensure data were only collected from those with the expected number of droplets, unambiguous color codes, and proper droplet merging to facilitate mixing of contents; 5) Align microwells of each given kChip across all imaged time points; and 6) Measure the average image intensity (reporter signal) from the GFP channel across the identified merged droplet area in each microwell.

Droplets were identified via a circular Hough transform applied to the summed dye channels. The average fluorescence intensities across dye channels were made into a 3D vector per droplet, which were projected into 2D space and clustered using the k-means clustering algorithm. Those clusters were mapped to the known kChip input barcodes, and droplets outside of a small radius around any given cluster centroid were dropped to ensure quality. Once identified and labeled, the droplets were assigned to wells and aligned across timepoints by cross-correlating the premerge image with a binary microwell mask template image and with the following postmerge timepoint image, respectively. Wells with incorrect numbers of assigned droplets were dropped from analysis. Merged droplets were segmented by a watershed algorithm within the 3D dye space, and any that showed signs of non-homogenous input mixing were also dropped.

For additional information on kChip image analysis, see ^40^.

### kChip data analysis

The activity from each *S. aureus* reporter in each microwell was calculated relative to the untreated controls:

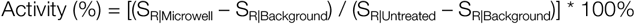

Where: S_R|Microwell_ = reporter signal for given microwell; S_R|Untreated_ = median reporter signal from microwells with only untreated control inputs; and S_R|Background_ = 10th percentile reporter signal from microwells with one or more antibiotic control inputs. (Note that for Screen 3, untreated controls were measured from *k* = 1 microwells on the kChips given the rare occurrences of *k* = 7 microwells that received seven untreated control inputs.)

The reporter activity for a given combination of inputs was calculated as the median activity for all microwells containing the given combination.

Analyses and visualizations for all screens were focused on wells with specific assemblages of inputs. In all screens, microwells that had been used for antibiotic or untreated control calculations were not included in downstream comparisons of biobank strain assemblage performances. Additionally, for Screen 1, analyses were focused on microwells that received either two distinct strains (AB) or two repeating stains (AA). Microwells receiving one strain and one medium-only control (A[]) were excluded from analyses and visualizations. For Screen 2, analyses were focused on microwells that received either three distinct strains selected from Screen 1 (ABC) or one strain and two medium-only controls (A[][]). All other assemblages (e.g., AAA, AAB, AA[], or AB[]) were excluded from analyses and visualizations. For Screen 3, analyses were focused on microwells that received seven community members (MMMMMMM), one Screen 2 selection and six community members (AMMMMMM), or three distinct Screen 2 selections and four community members (ABCMMMM). All other assemblages (e.g., ABMMMMM, ABCDMMM, AABMMMM, etc.) were excluded from analyses and visualizations.

The strength of observed suppression by each combination was categorized for each screen. In Screen 1, the combinations were categorized as exhibiting “strong suppression” (median relative *S. aureus* reporter activity <12.5%, or the equivalent of 3 doublings decrease, >8-fold decrease), “moderate suppression,” (≥12.5%; <“neutral”), or “neutral” effects. The “neutral” bound was determined based on the range of effects observed on the untreated control replicates: after calculating an individual effect size for each replicate, the bound was set at one standard deviation below the distribution mean. The lower bound for the P*agr* reporter was 84.8%, 71.9% for P*psmα*, 82.9% for P*sigB*, and 73.3% for P*gmk*. In Screen 2, the combinations were categorized as exhibiting “strong suppression” (<12.5%), “moderate suppression” (≥12.5%; <80%), or “neutral” effects (≥80%). In Screen 3, the combinations were categorized as exhibiting “strong suppression” (<12.5%) or “moderate suppression” (≥12.5%; <80%).

### Statistical analyses

In Screen 1, the effects of pairwise combinations of the biobank strains AB on the reporters were compared to those of their constituent strains AA and BB using paired t-tests with a Bonferroni-corrected significance threshold of 1.55 × 10^-6^. The microbe pairs were binned based on which of the paired differences were significant. For example, pairs were placed in the “AB<AA, AB” category when both the AB<AA and AB<BB comparisons were statistically significant; pairs were placed in the “AB>AA, AB” category when both the AB>AA and AB>BB comparisons were statistically significant. All other relationships were considered “neutral” and placed in the “AA≤AB≤BB” category; two examples of such distributions are depicted in Fig. 2f (AA≈AB≈BB and AA<AB<BB), but could also include (AA≈AB>BB) or (AA≈AB<BB).

The Screen 2 comparisons of ENS-002 performance to Strains N, T, and X were performed using two-sided Mann-Whitney U tests with a Bonferroni correction with a significance cut-off of *P* <0.017.

### Data visualization

kChip data analysis and visualization were performed using Python with pandas (v2.3.3), NumPy (v1.26.4), matplotlib (v3.10.7), and seaborn (v0.13.2). A maximum likelihood phylogenetic tree was constructed based on MUSCLE-aligned 16S sequences using Mega12^64^; the tree was visualized using FigTree (v1.4.5). Prism (v10.6.1) was used for microtiter and RHESA assay visualizations. Inkscape (v1.4.2) was used for figure organization.

### Off-kChip coculture assays

Pre-growth cultures of *S. aureus* reporter strains and treatment strains (ENS-002 or the BAA-1717 non-fluorescent WT strain) were normalized to the appropriate density and combined to achieve the desired starting densities for either a 1:1 ratio (2.5 × 10^7^ CFU/mL : 2.5 × 10^7^ CFU/mL) or a 10:1 ratio (2.5 × 10^8^ CFU/mL : 2.5 × 10^7^ CFU/mL) of *S. aureus* reporter : treatment strain or combination.

*S. aureus* reporter strains were normalized to 2X the desired final density in glucose-free TSB. Strains N, T, X, and non-fluorescent BAA-1717 cultures were first centrifuged at 3000 RCF for 10 min, and the resulting cell pellets were resuspended in phosphate-buffered saline (PBS). Then, these cultures were normalized to 2X the desired final density in PBS. To make the ENS-002 combination, Strains N, T, and X were combined in a 1:1:1 CFU ratio to achieve a total density of 2.5 × 10^7^ CFU/mL. Cultures were normalized using OD_600_ to CFU/mL conversion factors informed by a prior experiment (data not shown).

*S. aureus* cultures were then mixed in a 1:1 volume ratio with either an ENS-002 strain, the ENS-002 combination, or a PBS control in a total volume of 200 μL in triplicate in a black, clear bottom 96-well plate (Corning VV-01728-41). The 96-well plate was sealed with an optical seal (Millipore Sigma Z742420); holes were punched above each well for gas exchange. The plate was incubated in a BioTek Synergy H1 plate reader and grown at 37°C, with maximum-speed linear shaking. GFP fluorescence (Excitation: 485 nm, Emission: 511 nm, Gain: 50, Read Height: 7.5 mm) was measured every 7.5 minutes. After 24 hr, *S. aureus* CFU were enumerated.

### RHESA model

EpiDerm™ RHE models (EPI-200-EXP-AFAB, Version 3) were purchased from MatTek (Ashland, MA). Supplementary Fig. 8 provides a timeline of the RHESA protocol. Upon receipt, each 9 mm diameter model was transferred to a separate well of 6-well plates containing 1 mL of maintenance medium (EPI-100-NMM-HCF-AFAB). This medium does not contain antibiotics, antifungals, or hydrocortisone. Models were incubated aerobically at 37°C with 5% CO_2_. The models were kept in culture for approximately 20 h (a “pre-equilibration” period, which allows the skin models to recover from any stress caused by shipping). The maintenance medium was exchanged for fresh medium twice during the pre-equilibration period.

Following the pre-growth period, bacterial cultures were normalized using OD_600_ to CFU/mL conversion factors informed by a prior experiment (data not shown). At T = 0 hr, 1 x 10^6^ CFU *S. aureus* virulence reporter containing the P*agr*-*gfp* reporter construct was applied to the surfaces of the models in 150 μL of PBS. The models were then incubated aerobically at 37°C with 5% CO_2_ for 1 h to allow the *S. aureus* cells to settle onto the surfaces of the models. The PBS was then carefully aspirated without compromising the integrity of the models. At this point, the RHESA model was fully prepared for application of test materials.

At T = 1 hr, either 150 μL of PBS (“untreated”) or 1 x 10^6^ CFU of ENS-002 (∼3.3 x 10^5^ CFU of each strain) in 150 μL of PBS were applied to the surfaces of the appropriate models. The models were then incubated aerobically at 37°C with 5% CO_2_ for 1 h to allow the ENS-002 cells to settle onto the surfaces of the models. The PBS from the treatments was then carefully aspirated without compromising the integrity of the models. Models were then incubated aerobically at 37°C with 5% CO_2_ until T = 24 hr.

After the desired incubation period, the RHE models underwent an elution process to remove and enumerate the bacterial cells and quantify *S. aureus* virulence. A 6-mm biopsy punch was used to extract the center of each skin model. The punch was transferred to a tube containing an elution buffer (1X PBS, 0.05% Tween 20 (Sigma-Aldrich P9416), and 0.05% Triton X-100 (Sigma-Aldrich T8787)) and vortexed on maximum speed (3200 RPM) for 3 minutes. *S. aureus* CFU and total ENS-002 CFU from this solution were enumerated; the cell and population-level P*agr*-*gfp* fluorescence was measured by flow cytometry. Growth of each ENS-002 strain growth was not separately quantified in this experiment. In other experiments, Strain N, Strain T, and Strain X were separately applied to models and each observed to grow (data not shown).

### Flow cytometry

The samples eluted from each model were measured by flow cytometry to quantify GFP fluorescence from the P*agr*-*gfp* reporter. A set volume (13 μL) of each sample was run through an Attune CytPix Flow Cytometer at a rate of 12.5 μL/min. The BL1 detector (voltage: 370) for the blue laser was used for detecting GFP fluorescence (excitation: 488 nm, emission: 530 nm). The FSC voltage was set to 330; the SSC voltage was set to 410. An FSC threshold of 400 and an SSC threshold of 1800 were used. Flow Cytometry Standard files were exported from the Attune Cytometric Software (Version: 5.3.2415.0, Database Version: 4.3.002.4) for analyses. GFP height (H) values were used for all analyses. Fluorescent/virulent events were identified as those over a fluorescent cut-off (375 RFU) based on the bimodal distribution of fluorescence, where the GFP-positive peak was included above the threshold. From the fluorescent events, the total fluorescence (total virulence of the sample) was calculated as the sum of per-event fluorescent intensities within the gated population and median fluorescence (median virulence per cell) was calculated as the median fluorescence intensity within the gated population using a custom R script.

## Acknowledgements

The authors thank T. Scharschmidt (University of California, San Francisco), J. Sigler, M. Volpe, A. Sclafani, and A. Morton for helpful conversations through the discovery of ENS-002; J. Oh (Duke University) for discussions on RHESA model development; P. Blainey (Massachusetts Institute of Technology), C. Mitchell (Massachusetts General Hospital), D. West, and J. Koconis for comments on the manuscript; J. Merrick for figure design and formatting; O. Charlemagne for assistance in generating research cell banks; and R. Duquette for the procurement of laboratory reagents and supplies.

## Author Contributions

B.C., C.M.A.A., and J.K. conceptualized this work. B.C. and J.K. designed the biobanking effort. J.K. designed the kChip screens. B.C., A.H., and J.K. performed biobanking and kChip screens. K.A. performed all kChip image analysis and data analysis. K.A. and J.K. designed methods for selecting strains for continuation to subsequent screens. E.L.B. designed off-kChip experiments (microtiter plate and RHESA model experiments). E.L.B., O.S., H.A.A., and D.C. performed off-kChip experiments. E.A.Z. coordinated whole genome sequencing and antibiotic susceptibility assessments. A.L. generated ENS-002 cell banks. P.L. provided subject matter expertise. E.L.B., K.D.L., and J.K. wrote the paper.

## Additional Information

Supplementary Information is available for this paper.

Correspondence and requests for materials should be addressed to Jared Kehe.

## References

1. Kong, H. H. et al. Temporal shifts in the skin microbiome associated with disease flares and treatment in children with atopic dermatitis. Genome Res. 22, 850–859 (2012).

2. Meylan, P. et al. Skin Colonization by Staphylococcus aureus Precedes the Clinical Diagnosis of Atopic Dermatitis in Infancy. J. Invest. Dermatol. 137, 2497–2504 (2017).

3. Leyden, J. J., Marples, R. R. & Kligman, A. M. Staphylococcus aureus in the lesions of atopic dermatitis. Br. J. Dermatol. 90, 525–530 (1974).

4. Aly, R., Maibach, H. I. & Shinefield, H. R. Microbial Flora of Atopic Dermatitis. Arch. Dermatol. 113, 780–782 (1977).

5. Totté, J. E. E. et al. Prevalence and odds of Staphylococcus aureus carriage in atopic dermatitis: a systematic review and meta-analysis. Br. J. Dermatol. 175, 687–695 (2016).

6. Khadka, V. D. et al. The Skin Microbiome of Patients With Atopic Dermatitis Normalizes Gradually During Treatment. Front. Cell. Infect. Microbiol. 11, (2021).

7. Geoghegan, J. A., Irvine, A. D. & Foster, T. J. Staphylococcus aureus and Atopic Dermatitis: A Complex and Evolving Relationship. Trends Microbiol. 26, 484–497 (2018).

8. Kim, J., Kim, B. E., Ahn, K. & Leung, D. Y. M. Interactions Between Atopic Dermatitis and Staphylococcus aureus Infection: Clinical Implications. Allergy Asthma Immunol. Res. 11, 593–603 (2019).

9. Byrd, A. L., Belkaid, Y. & Segre, J. A. The human skin microbiome. Nat. Rev. Microbiol. 16, 143–155 (2018).

10. Huang, C. et al. Skin microbiota: pathogenic roles and implications in atopic dermatitis. Front. Cell. Infect. Microbiol. 14, 1518811 (2025).

11. Nakatsuji, T. & Gallo, R. L. The role of the skin microbiome in atopic dermatitis. Ann. Allergy Asthma Immunol. Off. Publ. Am. Coll. Allergy Asthma Immunol. 122, 263–269 (2019).

12. Jenul, C. & Horswill, A. R. Regulation of Staphylococcus aureus Virulence. Microbiol. Spectr. 7, 10.1128/microbiolspec.gpp3-0031–2018.

13. Cheung, G. Y. C., Bae, J. S. & Otto, M. Pathogenicity and virulence of Staphylococcus aureus. Virulence 12, 547–569 (2021).

14. Nakatsuji, T. et al. Competition between skin antimicrobial peptides and commensal bacteria in type 2 inflammation enables survival of S. aureus. Cell Rep. 42, (2023).

15. Ong, P. Y. et al. Endogenous Antimicrobial Peptides and Skin Infections in Atopic Dermatitis. N. Engl. J. Med. 347, 1151–1160 (2002).

16. Cho, S. H. et al. Preferential binding of Staphylococcus aureus to skin sites of Th2-mediated inflammation in a murine model. J. Invest. Dermatol. 116, 658–663 (2001).

17. Cho, S.-H., Strickland, I., Boguniewicz, M. & Leung, D. Y. M. Fibronectin and fibrinogen contribute to the enhanced binding of Staphylococcus aureus to atopic skin. J. Allergy Clin. Immunol. 108, 269–274 (2001).

18. Bitschar, K. et al. Staphylococcus aureus Skin Colonization Is Enhanced by the Interaction of Neutrophil Extracellular Traps with Keratinocytes. J. Invest. Dermatol. 140, 1054–1065.e4 (2020).

19. Arikawa, J. et al. Decreased Levels of Sphingosine, a Natural Antimicrobial Agent, may be Associated with Vulnerability of the Stratum Corneum from Patients with Atopic Dermatitis to Colonization by Staphylococcus aureus. J. Invest. Dermatol. 119, 433–439 (2002).

20. Simpson, E. L. et al. Rapid reduction in Staphylococcus aureus in atopic dermatitis subjects following dupilumab treatment. J. Allergy Clin. Immunol. 152, 1179–1195 (2023).

21. Leyden, J. J. & Kligman, A. M. The case for steroid—antibiotic combinations. Br. J. Dermatol. 96, 179–187 (1977).

22. Breuer, K., HÄussler, S., Kapp, A. & Werfel, T. Staphylococcus aureus: colonizing features and influence of an antibacterial treatment in adults with atopic dermatitis. Br. J. Dermatol. 147, 55–61 (2002).

23. Lever, R., Hadley, K., Downey, D. & Mackie, R. Staphylococcal colonization in atopic dermatitis and the effect of topical mupirocin therapy. Br. J. Dermatol. 119, 189–198 (1988).

24. Gong, J. Q. et al. Skin colonization by Staphylococcus aureus in patients with eczema and atopic dermatitis and relevant combined topical therapy: a double-blind multicentre randomized controlled trial. Br. J. Dermatol. 155, 680–687 (2006).

25. Hung, S.-H. et al. Staphylococcus colonization in atopic dermatitis treated with fluticasone or tacrolimus with or without antibiotics. Ann. Allergy. Asthma. Immunol. 98, 51–56 (2007).

26. Ewing, C. I. et al. Flucloxacillin in the treatment of atopic dermatitis. Br. J. Dermatol. 138, 1022–1029 (1998).

27. Harkins, C. P. et al. The widespread use of topical antimicrobials enriches for resistance in Staphylococcus aureus isolated from patients with atopic dermatitis. Br. J. Dermatol. 179, 951–958 (2018).

28. Schachner, L. A. et al. A Consensus on Staphylococcus aureus Exacerbated Atopic Dermatitis and the Need for a Novel Treatment. J. Drugs Dermatol. JDD 23, 825–832 (2024).

29. Lobefaro, F., Gualdi, G., Di Nuzzo, S. & Amerio, P. Atopic Dermatitis: Clinical Aspects and Unmet Needs. Biomedicines 10, 2927 (2022).

30. Ahuja, K., Sunkara, M. & Lio, P. Pathogenic Colonization: Defining the Role of Staphylococcus aureus in Atopic Dermatitis. Dermatitis® https://doi.org/10.1089/derm.2024.0401 (2025) doi:10.1089/derm.2024.0401.

31. Iwase, T. et al. Staphylococcus epidermidis Esp inhibits Staphylococcus aureus biofilm formation and nasal colonization. Nature 465, 346–349 (2010).

32. Nakatsuji, T. et al. Antimicrobials from human skin commensal bacteria protect against Staphylococcus aureus and are deficient in atopic dermatitis. Sci. Transl. Med. 9, eaah4680 (2017).

33. Myles, I. A. et al. Therapeutic responses to Roseomonas mucosa in atopic dermatitis may involve lipid-mediated TNF-related epithelial repair. Sci. Transl. Med. 12, eaaz8631 (2020).

34. Williams, M. R. et al. Quorum sensing between bacterial species on the skin protects against epidermal injury in atopic dermatitis. Sci. Transl. Med. 11, eaat8329 (2019).

35. Cogen, A. L. et al. Selective Antimicrobial Action Is Provided by Phenol-Soluble Modulins Derived from Staphylococcus epidermidis, a Normal Resident of the Skin. J. Invest. Dermatol. 130, 192–200 (2010).

36. Brown, M. M. et al. Novel Peptide from Commensal Staphylococcus simulans Blocks Methicillin-Resistant Staphylococcus aureus Quorum Sensing and Protects Host Skin from Damage. Antimicrob. Agents Chemother. 64, 10.1128/aac.00172-20 (2020).

37. Nakatsuji, T. et al. Development of a human skin commensal microbe for bacteriotherapy of atopic dermatitis and use in a phase 1 randomized clinical trial. Nat. Med. 27, 700–709 (2021).

38. Jacobson, M. E., Myles, I. A., Paller, A. S., Eichenfield, L. F. & Simpson, E. L. A Randomized, Double-Blind, Placebo-Controlled, Multicenter, 16-Week Trial to Evaluate the Efficacy and Safety of FB-401 in Children, Adolescents, and Adult Subjects (Ages 2 Years and Older) with Mild-to-Moderate Atopic Dermatitis. Dermatology 240, 85–94 (2023).

39. Myles, I. A. et al. First-in-human topical microbiome transplantation with Roseomonas mucosa for atopic dermatitis. JCI Insight 3, e120608, 120608 (2018).

40. Kehe, J. et al. Massively parallel screening of synthetic microbial communities. Proc. Natl. Acad. Sci. 116, 12804–12809 (2019).

41. Kulesa, A., Kehe, J., Hurtado, J. E., Tawde, P. & Blainey, P. C. Combinatorial drug discovery in nanoliter droplets. Proc. Natl. Acad. Sci. 115, 6685–6690 (2018).

42. Poudel, S. et al. Revealing 29 sets of independently modulated genes in Staphylococcus aureus, their regulators, and role in key physiological response. Proc. Natl. Acad. Sci. 117, 17228–17239 (2020).

43. Enright, M. C., Day, N. P., Davies, C. E., Peacock, S. J. & Spratt, B. G. Multilocus sequence typing for characterization of methicillin-resistant and methicillin-susceptible clones of Staphylococcus aureus. J. Clin. Microbiol. 38, 1008–1015 (2000).

44. Li, P. et al. Transcriptional Profiling of Staphylococcus aureus during the Transition from Asymptomatic Nasal Colonization to Skin Colonization/Infection in Patients with Atopic Dermatitis. Int. J. Mol. Sci. 25, 9165 (2024).

45. Key, F. M. et al. On-person adaptive evolution of Staphylococcus aureus during treatment for atopic dermatitis. Cell Host Microbe 31, 593–603.e7 (2023).

46. Piewngam, P. et al. Pathogen elimination by probiotic Bacillus via signalling interference. Nature 562, 532–537 (2018).

47. Wu, M.-Y. & Yao, X. Skin Microbiota and the Skin Barrier. Int. J. Dermatol. Venereol. 07, 18–26 (2024).

48. Linz, M. S., Mattappallil, A., Finkel, D. & Parker, D. Clinical Impact of Staphylococcus aureus Skin and Soft Tissue Infections. Antibiotics 12, 557 (2023).

49. Wang, X. et al. Staphylococcus aureus colonization and chronic hand eczema: a multicenter clinical trial. Arch. Dermatol. Res. 311, 513–518 (2019).

50. Kehe, J. et al. Positive interactions are common among culturable bacteria. Sci. Adv. 7, eabi7159 (2021).

51. Bawdon, D., Cox, D. S., Ashford, D., James, A. G. & Thomas, G. H. Identification of axillary Staphylococcus sp. involved in the production of the malodorous thioalcohol 3-methyl-3-sufanylhexan-1-ol. FEMS Microbiol. Lett. 362, fnv111 (2015).

52. Oh, J. & Voigt, A. Y. The human skin microbiome: from metagenomes to therapeutics. Nat. Rev. Microbiol. 23, 771–787 (2025).

53. Tao, R., Li, R. & Wang, R. Skin microbiome alterations in seborrheic dermatitis and dandruff: A systematic review. Exp. Dermatol. 30, 1546–1553 (2021).

54. Kurtz, S. et al. Versatile and open software for comparing large genomes. Genome Biol. 5, R12 (2004).

55. Chen, L. et al. VFDB: a reference database for bacterial virulence factors. Nucleic Acids Res. 33, D325–328 (2005).

56. Carattoli, A. et al. In Silico Detection and Typing of Plasmids using PlasmidFinder and Plasmid Multilocus Sequence Typing. Antimicrob. Agents Chemother. 58, 3895–3903 (2014).

57. Seidl, K. et al. Staphylococcus aureus CcpA affects virulence determinant production and antibiotic resistance. Antimicrob. Agents Chemother. 50, 1183–1194 (2006).

58. Tuchscherr, L. et al. Sigma Factor SigB Is Crucial to Mediate Staphylococcus aureus Adaptation during Chronic Infections. PLoS Pathog. 11, e1004870 (2015).

59. Horinouchi, S. & Weisblum, B. Nucleotide sequence and functional map of pC194, a plasmid that specifies inducible chloramphenicol resistance. J. Bacteriol. 150, 815–825 (1982).

60. Lee, M. E., DeLoache, W. C., Cervantes, B. & Dueber, J. E. A Highly Characterized Yeast Toolkit for Modular, Multipart Assembly. ACS Synth. Biol. 4, 975–986 (2015).

61. Gibson, D. G. et al. Enzymatic assembly of DNA molecules up to several hundred kilobases. Nat. Methods 6, 343–345 (2009).

62. Engler, C., Kandzia, R. & Marillonnet, S. A one pot, one step, precision cloning method with high throughput capability. PloS One 3, e3647 (2008).

63. Bose, J. L. The Genetic Manipulation of Staphylococci: Methods and Protocols. (Humana Press).

64. Kumar, S. et al. MEGA12: Molecular Evolutionary Genetic Analysis Version 12 for Adaptive and Green Computing. Mol. Biol. Evol. 41, msae263 (2024).

