## Supplementary Tables and Figures for "Systematic discovery of a topical bacterial consortium that targets *Staphylococcus aureus* to treat atopic dermatitis"

### Supplementary Table and Figure Legends

**Supplementary Table 1 | Genus representation across discovery process**
[a] Genus assignments were based on the top BLAST hit for the 16S sequences of each strain. A quality cut-off of 750 nt and >97% identity was used.
[b] Five biobank isolates were found to be impure, mixed cultures (contained two or more distinct species).
[c] Three strains failed 16S sequencing in two separate attempts. Because they were not included in the screen or in downstream processes, they were not re-attempted.

**Supplementary Table 2. *S. aureus* reporters**
[a] The reporters were constructed in an entry vector (Methods). Promoter regions were amplified from the *S. aureus* strain ATCC 35556 (SA113). Most reporter strains were generated by transforming the resulting plasmids into the ATCC BAA-1717 background. Five additional *S. aureus* strains, ATCC 12600, ATCC 43300, ATCC 53657, BAA-39, and BAA-2094, were transformed with the P*gmk-gfp* reporter plasmid.

**Supplementary Fig. 1 | Screen 1 P*psma* heat map** The median effect of each pairwise biobank combination on *S. aureus* P*psma-gfp* activity relative to the untreated control. A maximum likelihood phylogenetic tree was constructed based on 16S sequences to sort individual strains (Methods). Two of the strains that were screened were later found to be polymicrobial and were not included in the tree. Colored boxes indicate genera of the top 28 strain selections resulting from this screen. Gray boxes capture all remaining genera in Supplementary Table 1 that were not included in the top 28. Arrows on the right-hand side of the heat map indicate the 28 strains that were selected for further testing; "*" indicates the eventually-selected ENS-002 strains.Data shown are median effects from a median replicate count of 32. AFU, arbitrary fluorescence units.

**Supplementary Fig. 2 | Screen 1 P*gmk* heat map** The median effect of each pairwise biobank combination on *S. aureus* P*gmk-gfp* activity relative to the untreated control. A maximum likelihood phylogenetic tree was constructed based on 16S sequences to sort individual strains (Methods). Two of the strains that were screened were later found to be polymicrobial and were not included in the tree. Colored boxes indicate genera of the top 28 strain selections resulting from this screen. Gray boxes capture all remaining genera in Supplementary Table 1 that were not included in the top 28. Arrows on the right-hand side of the heat map indicate the 28 strains that were selected for further testing; "*" indicates the eventually-selected ENS-002 strains. Data shown are median effects from a median replicate count of 33. AFU, arbitrary fluorescence units.

**Supplementary Fig. 3 | Screen 1 P*sigB* heat map** The median effect of each pairwise biobank combination on *S. aureus* P*sigB-gfp* activity relative to the untreated control. A maximum likelihood phylogenetic tree was constructed based on 16S sequences to sort individual strains (Methods). Two of the strains that were screened were later found to be polymicrobial and were not included in the tree. Colored boxes indicate genera of the top 28 strain selections resulting from this screen. Gray boxes capture all remaining genera in Supplementary Table 1 that were not included in the top 28. Arrows on the right-hand side of the heat map indicate the 28 strains that were selected for further testing; "*" indicates the eventually-selected ENS-002 strains. Data shown are median effects from a median replicate count of 36. AFU, arbitrary fluorescence units.

**Supplementary Fig. 4 | Screen 1 cross-reporter scatter plots a-d**, The median effect of each biobank strain AA (**a**,**c**) or pairwise combination AB (**b**,**d**) on either P*psma-gfp* and P*gmk-gfp* (**a**,**b**), or P*sigB-gfp* and P*gmk-gfp* (**c**,**d**) activity relative to the untreated control. Blue triangles indicate the strains or combinations containing two inputs of the 28 strains selected for advancement; gray circles indicate all remaining data. Dashed horizontal and vertical lines indicate bounds for strong (<12.5%) and moderate suppression (≥12.5%, <neutral). Data shown are median effects from a median replicate count of 32 (P*psma*), 33 (P*gmk*), and 36 (P*sigB*). AFU, arbitrary fluorescence units.

**Supplementary Fig. 5 | Screen 2 cross-reporter scatter plots a-e**, The median effect of all non-repeating three-wise combinations of biobank strains, excluding those microwells that received any medium-only inputs on P*gmk-gfp* and P*psmα-gfp* (**a**), P*saeR-gfp* (**b**), P*sigB-gfp* (**c**), P*ccpA-gfp* (**d**) activity relative to the untreated control. Yellow circles indicate the combinations that received three non-repeating inputs of 14 strains selected for advancement; blue triangles indicate all remaining data. The ENS-002 combination is indicated with the black circle. Dashed horizontal and vertical lines indicate bounds for strong (<12.5%) and moderate suppression (≥12.5%, <80%). Data shown are median effects from a median replicate count of 37 (P*sigB*), 31 (P*psmα*), 35 (P*saeR*), 40 (P*ccpA*), and 35 (P*gmk*). AFU, arbitrary fluorescence units.

**Supplementary Fig. 6 | Pie chart of community members** Pie chart detailing the genus representation of the 78 community members used in Screen 3.

**Supplementary Fig. 7 | ENS-002 individual strain performance in Screen 3 a,** A schematic depicting the types of seven-wise combinations of interest in (**b-d**) that were generated from the 14 strain selections from Screen 2 (See Fig. 1b) (yellow highlighting; dark blue highlighting for ENS-002) and 78 community members (see Supplementary Fig. 6) (gray dashed circles). **b**-**d**, For the *S. aureus* reporters P*agr-gfp* (**b**), P*psmα-gfp* (**c**), and P*gmk-gfp* (**d**), two distributions of *S. aureus* activity relative to the untreated control are shown. The background lightest gray distribution shows the relative *S. aureus* activity of all assembled MMMMMMM combinations (counts on right axes). In the case a particular community was assembled more than once, the median is reported. The forefront distributions show the relative *S. aureus* activity of all assembled AMMMMMM combinations (counts on left axes) [dark gray for strong suppression (<12.5%); medium gray for moderate suppression (≥12.5%, <80%); light gray for neutral (≥80%)]. The blue circles along the X-axes indicate the median reporter activity observed in microwells receiving one input of Strain N (star), Strain T (triangle), or Strain X (square). For the P*agr*, P*psmα*, and P*gmk* reporters, data were collected from 1618, 1760, 1990 MMMMMMM microwells and 4129, 5011, 5345 AMMMMMM microwells, respectively. A median of 293, 345, and 380 replicates were collected for the P*agr*, P*psmα*, and P*gmk* reporters, respectively. AFU, arbitrary fluorescence units.

**Supplementary Fig. 8 | RHESA timeline** A schematic illustrating the timing of the RHESA (Reconstructed Human Epidermidis *S. aureus* Activity) model (Using RHE models: EpiDerm™ EPI-200-EXP-AFAB, Version 3, purchased from MatTek in Ashland, MA). Each tick on the timeline represents 1 hour, where *S. aureus* is applied at T = 0 h. Baseline levels of *S. aureus* and ENS-002 were enumerated at T = 2 h.

**Supplementary Fig. 9 | Fluorescence distributions from the RHESA model eluted samples** Histograms of fluorescent events (GFP) detected by running a set volume of the eluted samples from the PBS-treated (**a**) and ENS-002-treated (**b**) RHESA models shown in Fig. 6 through an Attune CytPix Flow Cytometer at T = 24 h. Overlapping distributions are shown for the three replicates for each condition. Values <375 AFU were categorized as “not fluorescent” and represent non-fluorescent *S. aureus* cells, ENS-002 cells, or debris (represented by gray block). These values were excluded from summary analyses. The median fluorescence values reported in Fig. 6 indicate the median value of the fluorescent events (>375 AFU) and are indicated here by the green and blue circles on the X-axes. AFU, arbitrary fluorescence units.

### SUPPLEMENTARY TABLE 1

| Genus[a] | Count in Biobank of 609 Isolates | Count in Screen 1 Input List | Count in Screen 1 Selection List | Count in Screen 2 Selection List | Count in Screen 3 Community List |
|---|---|---|---|---|---|
| *Alkalihalobacillus* | 1 | 1 | 0 | 0 | 0 |
| *Aneurinibacillus* | 10 | 4 | 0 | 0 | 2 |
| *Arsenicicoccus* | 1 | 1 | 0 | 0 | 0 |
| *Aureimonas* | 1 | 1 | 0 | 0 | 0 |
| *Bacillus* | 61 | 19 | 9 | 8 | 4 |
| *Brevibacillus* | 15 | 9 | 1 | 0 | 1 |
| *Calidifontibacter* | 1 | 1 | 0 | 0 | 0 |
| *Cellulomonas* | 1 | 1 | 0 | 0 | 0 |
| *Corynebacterium* | 3 | 2 | 0 | 0 | 0 |
| *Cytobacillus* | 3 | 2 | 0 | 0 | 0 |
| *Dermabacter* | 4 | 2 | 0 | 0 | 1 |
| *Dermacoccus* | 24 | 8 | 1 | 0 | 4 |
| *Heyndrickxia* | 1 | 1 | 0 | 0 | 0 |
| *Janibacter* | 2 | 1 | 0 | 0 | 0 |
| *Kocuria* | 6 | 3 | 1 | 1 | 0 |
| *Lysinibacillus* | 6 | 3 | 2 | 1 | 1 |
| *Macrococcus* | 1 | 1 | 1 | 0 | 0 |
| *Microbacterium* | 5 | 3 | 0 | 0 | 0 |
| *Micrococcus* | 42 | 9 | 1 | 0 | 2 |
| *Neisseria* | 1 | 1 | 0 | 0 | 0 |
| *Oceanobacillus* | 2 | 1 | 0 | 0 | 1 |
| *Paenibacillus* | 20 | 8 | 1 | 0 | 1 |
| *Priestia* | 1 | 0 | 0 | 0 | 0 |
| *Pseudomonas* | 5 | 2 | 0 | 0 | 0 |
| *Psychrobacillus* | 2 | 1 | 0 | 0 | 1 |
| *Rhodococcus* | 1 | 0 | 0 | 0 | 0 |
| *Roseomonas* | 4 | 0 | 0 | 0 | 3 |
| *Rothia* | 1 | 1 | 0 | 0 | 0 |
| *Siminovitchia* | 2 | 0 | 0 | 0 | 0 |
| *Solibacillus* | 1 | 1 | 1 | 1 | 0 |
| *Sporosarcina* | 5 | 2 | 0 | 0 | 1 |
| *Staphylococcus* | 347 | 84 | 10 | 3 | 55 |
| *Streptococcus* | 20 | 5 | 0 | 0 | 0 |
| *Thermocrispum* | 1 | 0 | 0 | 0 | 0 |
| Mixed[b] | 5 | 2 | 0 | 0 | 1 |
| Unclassified[c] | 3 | 0 | 0 | 0 | 0 |
| **Total** | **609** | **180** | **28** | **14** | **78** |

### SUPPLEMENTARY TABLE 2

| Reporter Category | Reporter Construct[a] | Boundaries of promoter |
|---|---|---|
| Virulence – Quorum Sensing | P*agr-gfp* | The promoter region upstream of *agrB* (extending 184 bp upstream of the *agrB* start codon through the first 58 bp of *agrB*) |
| Virulence – Toxin Production | P*psmα-gfp* | The promoter region upstream of *psmα-1* (the 267 bp immediately upstream of the *psmα-1* start codon) |
| Virulence – Regulation | P*saeR-gfp* | The promoter region upstream of *saeR* (the 748 bp immediately upstream of the *saeR* start codon) |
| Metabolic Activity | P*gmk-gfp* | The promoter region upstream of *gmk* (the 300 bp immediately upstream of the *gmk* start codon) |
| Metabolic Activity | P*ccpA-gfp* | The promoter region of *ccpA* (the 500 bp immediately upstream of the *ccpA* start codon) |
| Stress | P*sigB-gfp* | The promoter region upstream of *rsbV* (the 300 bp immediately upstream of the *rsbV* start codon; this promoter regulates an operon containing *rsbV*, *rsbW*, and *sigB*) |

### SUPPLEMENTARY FIGURE 1

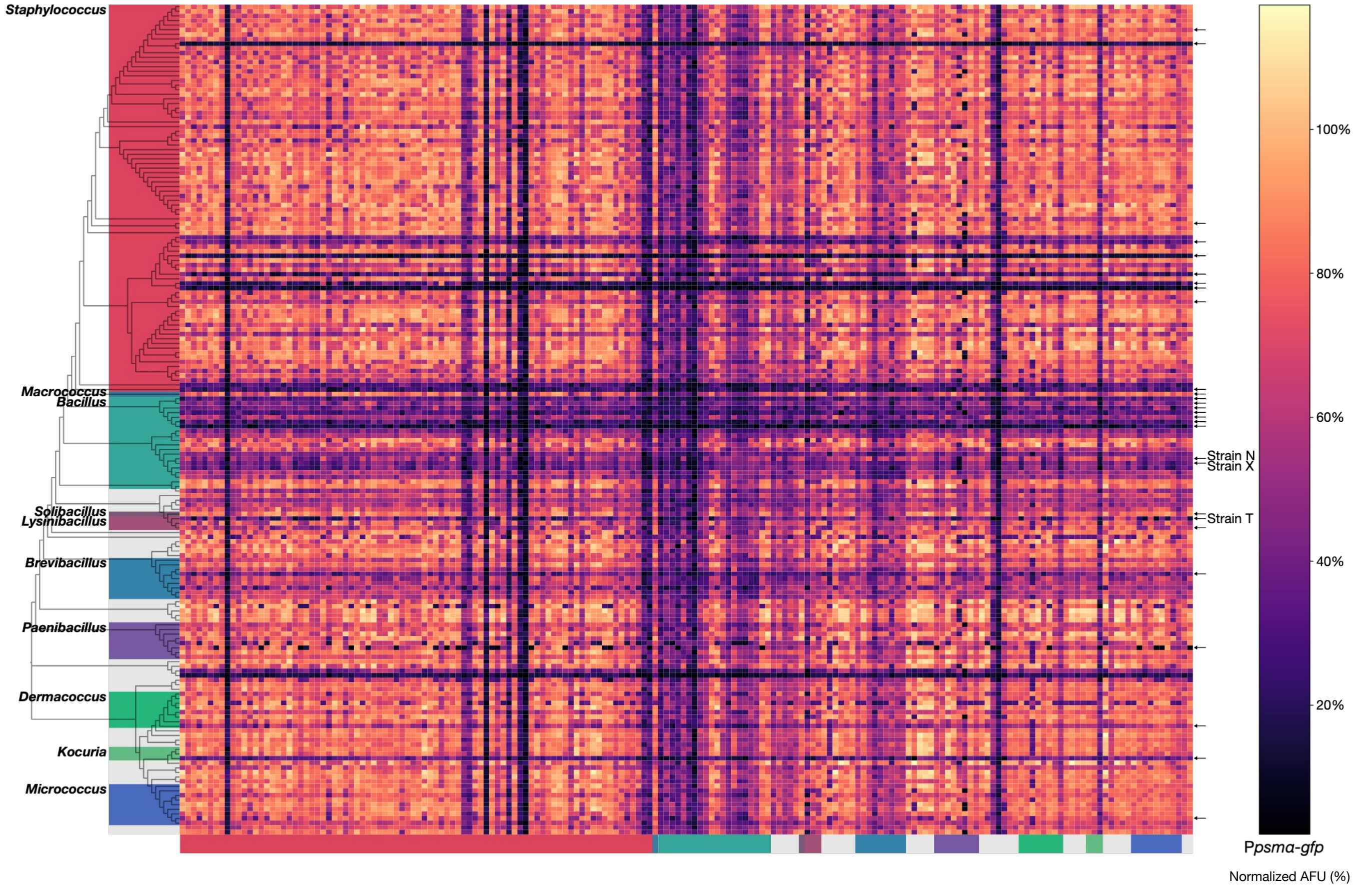

### SUPPLEMENTARY FIGURE 2

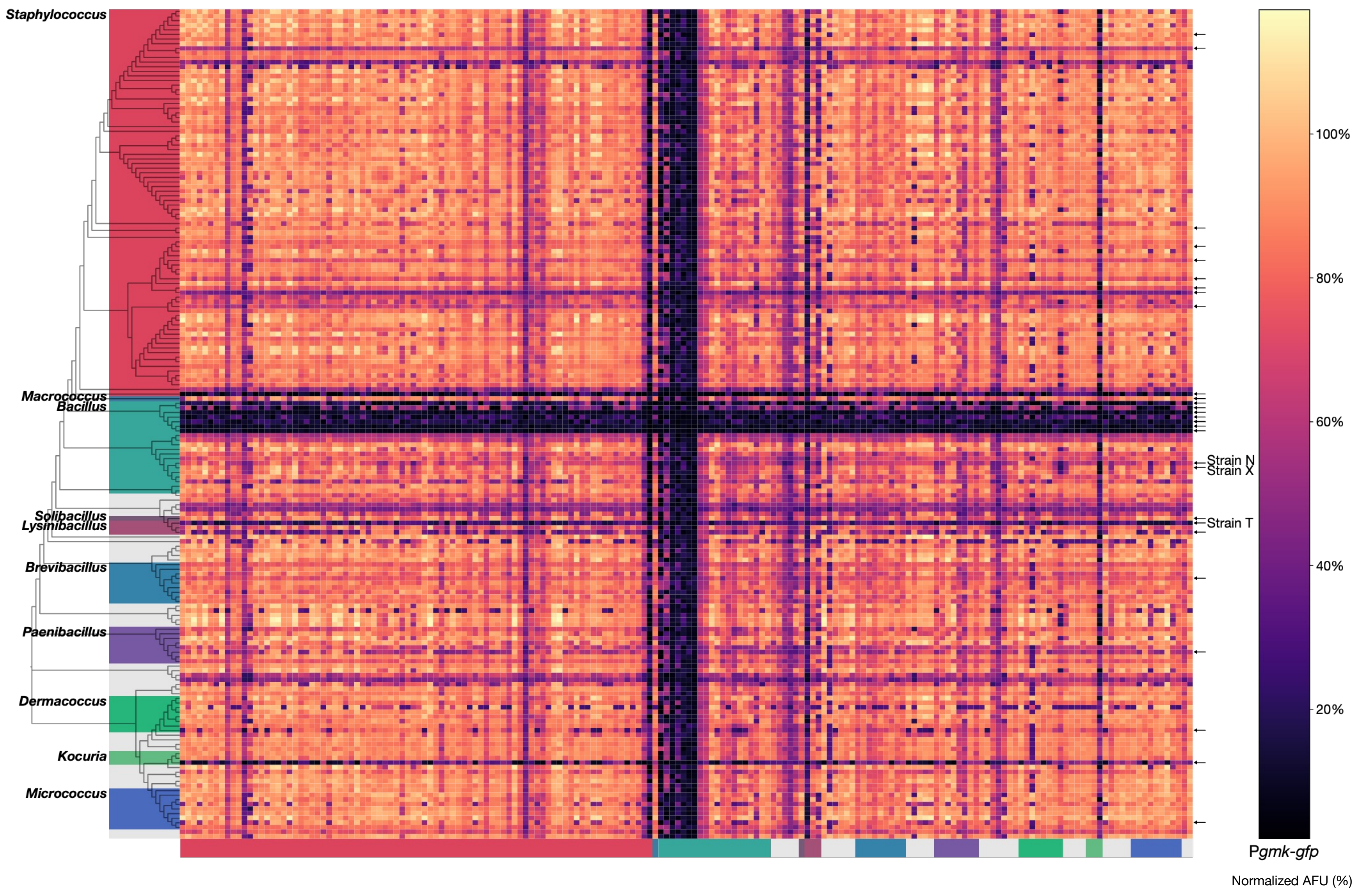

### SUPPLEMENTARY FIGURE 3

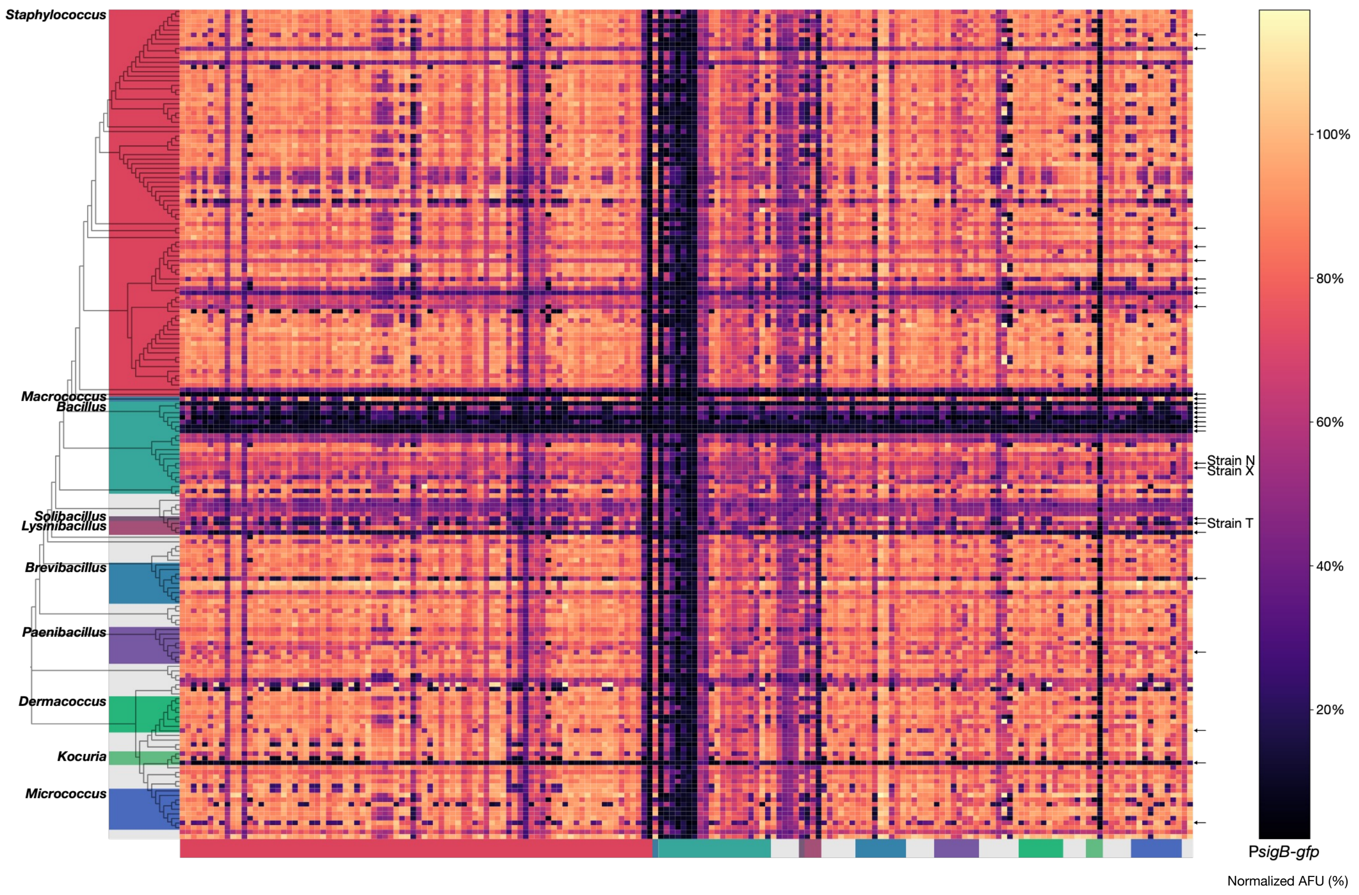

### SUPPLEMENTARY FIGURE 4

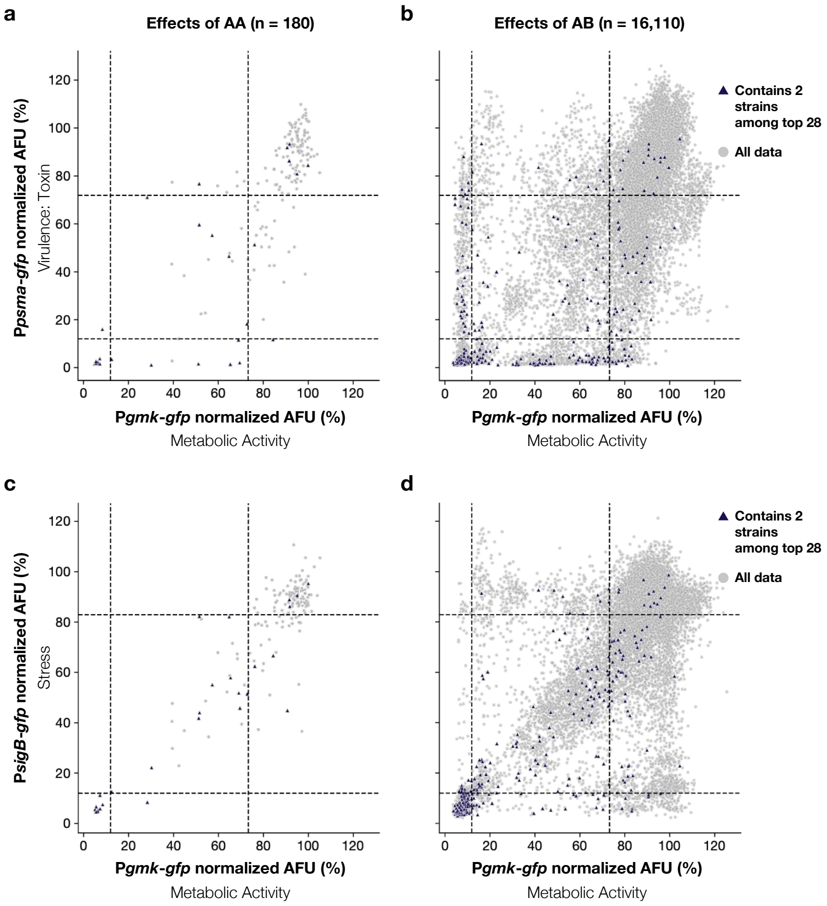

### SUPPLEMENTARY FIGURE 5

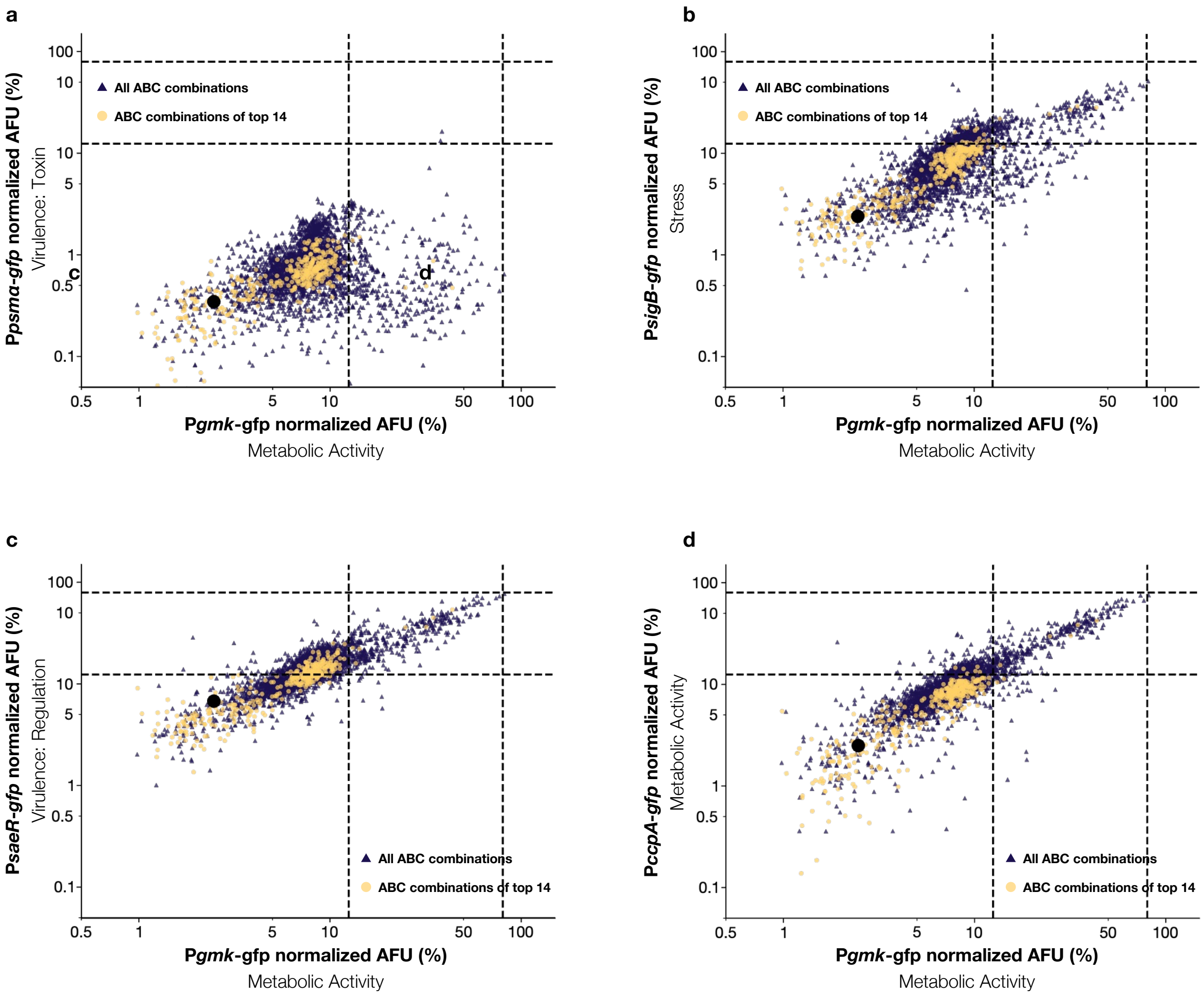

### SUPPLEMENTARY FIGURE 6

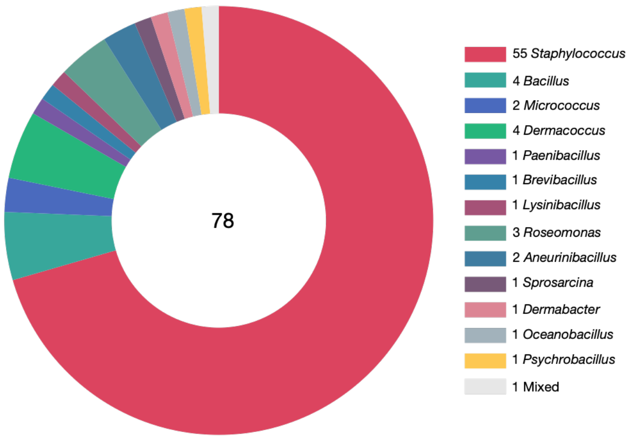

### SUPPLEMENTARY FIGURE 7

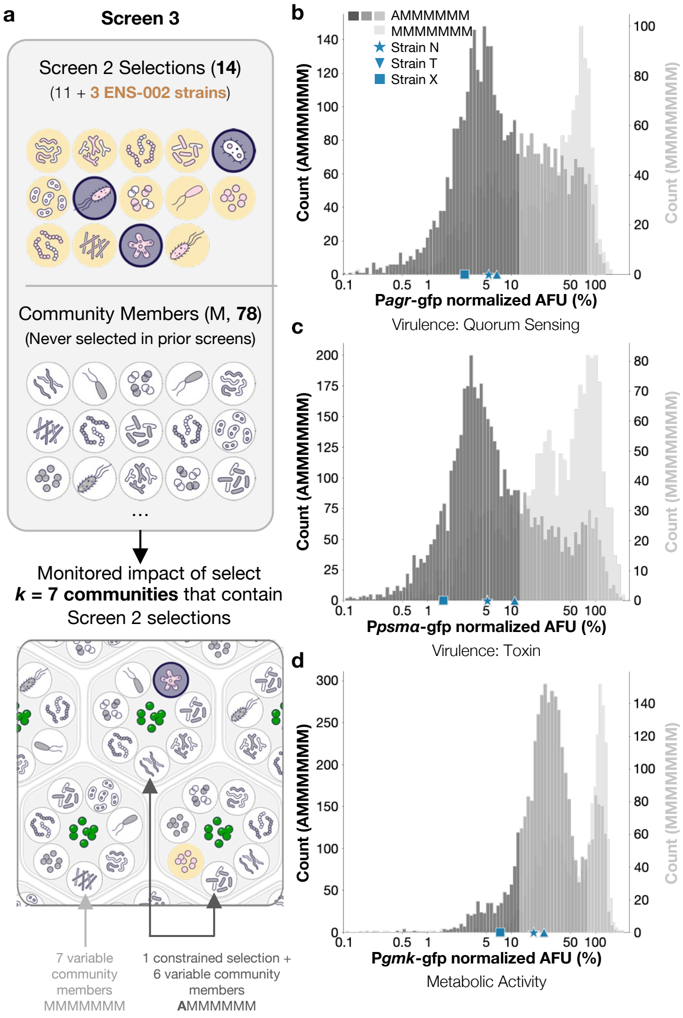

### SUPPLEMENTARY FIGURE 8

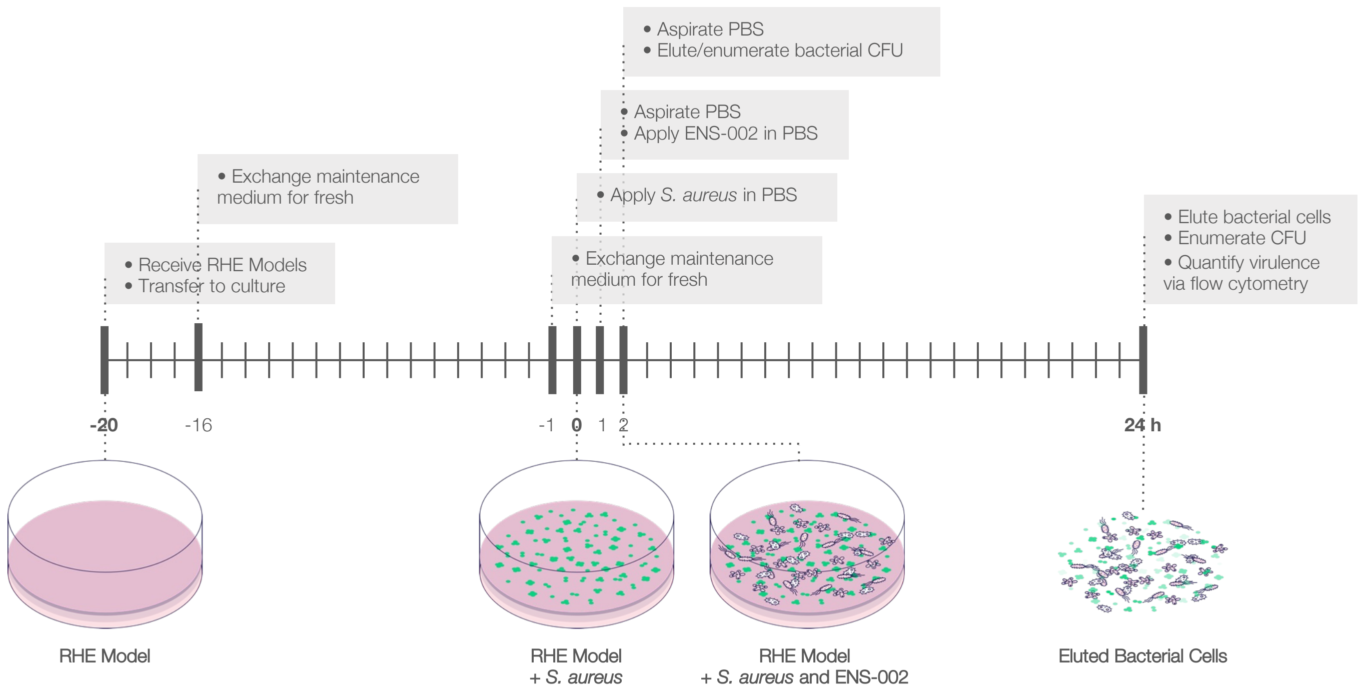

### SUPPLEMENTARY FIGURE 9

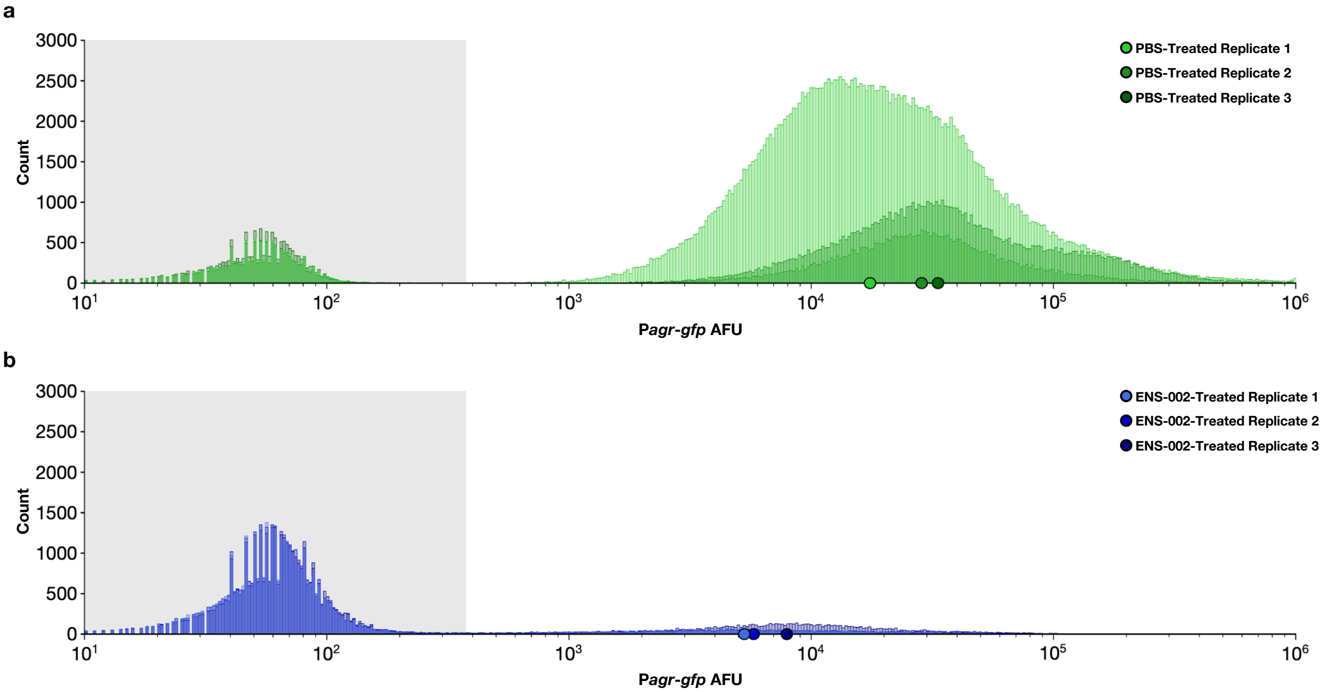
